# Construction of an engineered *Escherichia coli* with enhanced intestinal colonization and anti-inflammatory efficacy in colitis

**DOI:** 10.1101/2025.08.01.668021

**Authors:** Peijun Yu, Wenjing Zhou, Chunyang Li, Qiang Sun, Yunpeng Yang

## Abstract

Engineered probiotics are considered as effective and safe therapeutic strategies for the treatment of various diseases. *Escherichia coli* Nissle 1917 (EcN) has been widely used as a chassis strain for its safety and well-established genetic manipulation system. The limited intestinal colonization ability of EcN hampers its potential as a chassis for constructing synthetic probiotics in therapeutic applications. Here, an engineered EcN strain with high gastric acid and bile salts tolerance as well as enhanced intestinal adhesion ability was constructed by improving the expression of gastric acid and bile salts tolerance-associated genes and strengthening the expression of curli fiber formation genes, respectively. Meanwhile, oral administration of this engineered strain to colitis mice alleviated the disease severity and restored the disordered gut microbiome by decreasing the abundance of *Escherichia-Shigella*, whereas increasing *norank_f_Muribaculaceae*. We engineered EcN to improve its probiotic properties and anti-colitis efficacy, thereby establishing a platform for precision-designed synthetic probiotics.

## Introduction

Gut microbiota, the complex and dynamic ecosystem containing bacteria, viruses, fungi, yeasts, and other single-cell organisms, resides in the intestinal tract of the host and plays pivotal roles in maintaining the health state of host by affecting the nutrient metabolism, immune system, and neurological behavior [1]. The dysbiosis of gut microbiome is closely associated with various diseases, such as diabetes, obesity, hypertensive heart disease, inflammatory bowel disease, and neurological disorders [2]. Thus, reshaping the disordered gut microbiome is regarded as a potential strategy for disease treatment and attracts more and more attentions.

As reported, the administration of probiotics can benefit the host by modulating the gut microbial composition and inhibiting the intestinal colonization of various pathogens [3]. Among these probiotics, EcN, a non-pathogenic *E. coli* strain, benefits hosts by producing antimicrobial peptides to inhibit pathogens, secreting β-defensins to strengthen the intestinal barrier, and modulating immune responses through host interactions [4, 5]. Since the mature genetic engineering and synthetic biology toolkit had been developed in EcN, it had been engineered to treat a spectrum of diseases, e.g., tumors [6, 7], intestinal colitis [8, 9], and neurological disorders [10]. However, the poor intestinal colonization ability of EcN limits the therapeutic application of EcN-derived synthetic probiotics.

Until now, various strategies have been employed to enhance the intestinal colonization ability of EcN, such as facilitating the synthesis of bioactive metabolites that effectively suppress the growth of pathogens while promoting an optimized microenvironment for the growth of itself [11]; Enhancing the intestinal competitiveness of EcN by improving its nutrients (e.g., tripeptides) utilization ability [12]. Apart from the strategies mentioned above, EcN had been encapsulated with protective and adhesive materials to improve its tolerance towards the harsh gastrointestinal (GI) environment (e.g., gastric acid, bile salts, digestive enzymes, and intestinal motility) [13, 14]. However, constrained by the non-renewable of the protective and adhesive materials and the time-consuming procedure of encapsulation, new strategies should be developed to enhance the resistance of EcN toward harsh GI conditions.

In this study, we tried to improve the probiotic properties of EcN by applying two strategies: 1) Co-expressing the acid tolerance activator PatE and bile acid efflux transporter MdtM to counteract the gastric acid and bile acid stress; 2) Enhancing the intestinal adhesion of EcN by strengthening the expression of curli fiber formation genes, thus prolonging its intestinal retention. By integrating these functional elements into the genome of EcN, we constructed an engineered probiotic Syn3, whose intestinal colonization ability was superior than that of EcN. Oral administration of Syn3 to the dextran sulfate sodium (DSS)-induced colitis mice alleviated the disease severity by increasing the cecum and colon length, reducing the intestinal permeability and pro-inflammatory cytokines, and restoring the disordered gut microbiome by decreasing the abundance of *Escherichia-Shigella*, whereas increasing *norank_f_Muribaculaceae*. Collectively, our study constructed an engineered strain with improved probiotic properties and enhanced anti-colitis efficacy (Fig.1), which may serve as an excellent and stable chassis for the construction of EcN-derived synthetic probiotics.

**Fig. 1.**
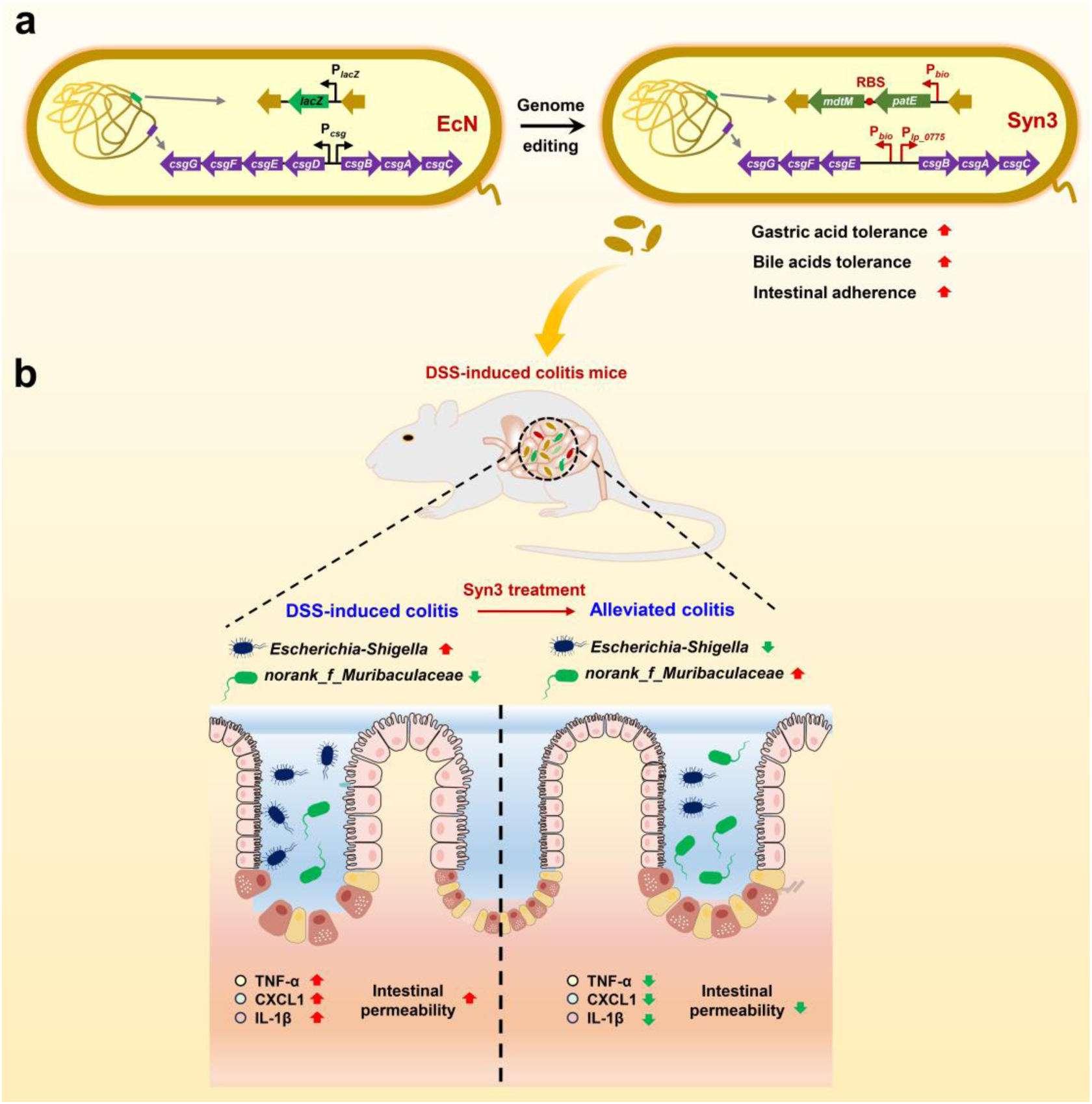
Oral administration of Syn3, an engineered EcN strain with enhanced probiotic properties, alleviates the disease phenotype of DSS-induced colitis mice. a EcN was genetically engineered to overexpress PatE and MdtM as well as strengthen the production of curli fibers by replacing the promoter of Csg operon with two strong promoters P*_lp_0775_* and P*_bio_*. EcN, *Escherichia coli* Nissle 1917; Syn3, engineered EcN strain. b Syn3 administration significantly alleviates the disease phenotype of DSS-induced colitis mice by reducing intestinal inflammation, protecting the intestinal barrier, and regulating the intestinal microbiome.

## Results

### Improving the gastric acid and bile salts tolerance of EcN by overexpressing PatE and MdtM

Gastric acid and bile acids are critical barriers that impede the intestinal colonization of gut microbes [15, 16]. To overcome these challenges, we tried to overexpress the gastric acid and bile acids tolerance-associated genes in EcN. Firstly, we screened the strong promoter that suitable for EcN by using the LacZ-based reporter system. The highest level of β-galactosidase activity was observed when it was under the control of P*_bio_*, the promoter that derived from the biotin synthesis gene cluster of *Clostridium acetobutylicum* (Supplementary Fig. S1a, b). Thus, this promoter was used to enhance the expression of functional genes in EcN.

As reported, *E. coli* can resist acidic stress via five acid stress response (AR1-5) systems [17]. Among these systems, the structural components and detailed acid-resistant mechanism of AR1 are poorly understood. Here, the specific mechanisms of AR2-5 are illustrated in EcN (Fig. 2a). As for the acid tolerance-related regulators, PatE (CIW80_12240), a homolog of putative AraC-like regulatory protein, had been proved to be an activator for the acid resistance-associated genes of Enterohemorrhagic *Escherchia coli* [18]. Thus, we overexpressed PatE in EcN under the control of P*_bio_*, yielding the engineered strain EcN-PatE. The EcN strain containing the empty vector (EcN-P) was used as control. The growth curve of EcN-PatE was comparable to that of EcN-P, indicating that overexpression of PatE did not impose a growth burden on EcN (Supplementary Fig. S2a). As reported, the AR1-3 pathways can be activated by low pH and enable bacteria to survive in extremely acidic environment (pH 2.5), whereas the AR4 and AR5 pathways allow bacteria to survive in moderately acidic environment (pH 4.5) [19]. Thus, we investigated the AR1-3 acid tolerance system-based survival rate of EcN-P and EcN-PatE under the extreme acid condition (pH = 2.5). As shown in Fig. 2b, the AR1-3 systems could be activated by PatE and were responsible for the higher acid survival rate of EcN-PatE. Furthermore, we compared the expression level of acid tolerance-associated genes between EcN-P and EcN-PatE, and found that most of the genes belonging to AR2-5 acid tolerance systems were up-regulated in EcN-PatE (Fig. 2c).

**Fig. 2.**
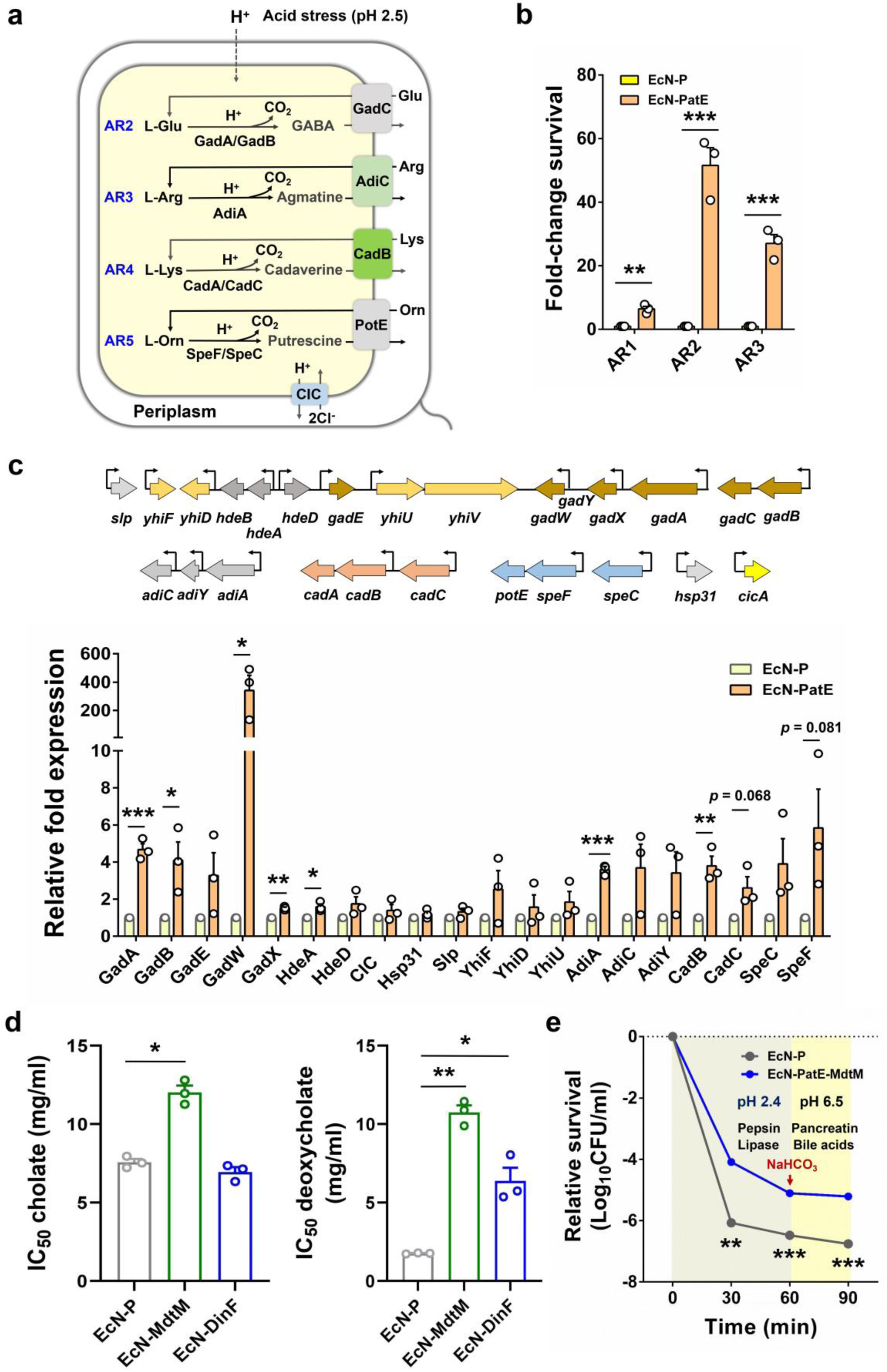
Overexpression of PatE and MdtM improved the gastric acid and bile salts tolerance of EcN. **a** Schematic illustration of acid response systems (AR2-AR5) in EcN. **b** The AR1-3 acid tolerance system-based survival rate of EcN-P (control strain) and EcN-PatE (PatE overexpression strain) under the extreme acid condition (pH = 2.5). **c** qRT-PCR analysis for the expression of acid tolerance-related genes in EcN-P and EcN-PatE. **d** The cholate and deoxycholate tolerance ability of EcN-P, EcN-MdtM (MdtM overexpression strain) and EcN-DinF (DinF overexpression strain). The concentration of bile salts that inhibit the growth of bacterial cell by 50% (IC_50_) was used to quantify bile salts tolerance of engineered strains. **e** Relative survival of EcN-P and EcN-PatE-MdtM (PatE and MdtM overexpression strain) at 0, 30, 60 and 90 minutes under the gastrointestinal-tract mimicking environment. The data are presented as mean ± SEM. The statistical significance between two groups were analyzed using *t*-test. The statistical significance between different groups (more than 3 groups) was calculated by using one-way analysis of variance (ANOVA) (\**P* < 0.05, \*\**P* < 0.01, \*\*\**P* < 0.001).

Following the gastric acid challenge, the next obstacle that restrict the intestinal colonization of EcN is bile salts in small intestine. As the potent detergents and oxidizing agents, bile salts can inhibit the survival of gut microbes by disrupting their extracellular and intracellular membranes [20]. Among the bile salts components, sodium cholate and sodium deoxycholate accounted for a main proportion. Here, we tried to improve the cholate and deoxycholate tolerance of EcN by overexpressing the bile salts tolerance-associated protein. To fulfill this goal, MdtM (CIW80_17355) and DinF (CIW80_15360), the two bile salts exportation-associated multidrug resistance transporters [21, 22], were overexpressed in EcN, yielding the engineered strain EcN-MdtM and EcN-DinF. The growth of EcN-MdtM and EcN-DinF were comparable to that of EcN-P (Supplementary Fig. S2b). Then, the bile salts tolerance of these two engineered strains was assessed by inoculating the grown cells of EcN-P, EcN-MdtM, and EcN-DinF into the LB liquid medium containing various concentrations of cholate (8, 12, 16, 24, 32, and 40 mg/ml) and deoxycholate (4, 8, 12, 24, 32, 48, and 64 mg/ml). Here, the concentration of bile salts that inhibited the growth of bacteria by 50% (IC_50_) was used to quantify the bile salts tolerance ability of bacteria. As shown in Fig. 2d, the cholate and deoxycholate tolerance ability of EcN-MdtM were superior to that of EcN-P and EcN-DinF. EcN-DinF can only resist the deoxycholate. Thus, MdtM was proved to be the functional bile salts exporter.

To improve the gastric acid and bile salts tolerance of EcN simultaneously, PatE and MdtM were co-expressed in EcN, yielding the engineered strain EcN-PatE-MdtM. The gastric acid and bile salts tolerance of EcN-PatE-MdtM and EcN-P were evaluated by incubating these microbes into the gastrointestinal-tract mimicking system [23]. In this experiment, the bacterial samples were collected at 0, 30, 60, and 90 minutes, serial diluted, and spread on the ampicillin-containing LB plates. The colony-forming unit (CFU) of EcN-PatE-MdtM and EcN-P were counted and used to calculate the relative survival rate of these two bacteria at specific time points according to the following equation: survival rate=CFU (30, 60, and 90 minutes)/CFU (0 minutes). As shown in Fig. 2e, the survival rate of EcN-PatE-MdtM was higher than that of EcN-P.

All these results indicated that PatE and MdtM are two functional genes that contribute to the gastric acid and bile salts tolerance of EcN, respectively.

### Enhancement of EcN adhesion to intestinal epithelial cells via strengthened curli amyloid fiber formation

The intestinal epithelial cell adhesion ability of gut microbes determines their intestinal colonization [24]. As reported, overexpression of amyloid curli fibers can facilitate the adherence of EcN to the intestinal mucosa, thereby improving its intestinal colonization ability [9]. In *E. coli*, the two divergently transcribed operons *csgBAC* and *csgDEFG* are responsible for the formation of extracellular amyloid fiber [25]. Thus, we tried to enhance the curli fibers formation of EcN by replacing the natural weak promoters of *csgBAC* and *csgDEFG* operons with P*_lp_0775_* and P*_bio_*, the two strong promoters that we previously screened in EcN (Fig. 3a and Supplementary Fig. S1). Besides, the curli fiber formation-associated transcriptional regulator CsgD was deleted from the genome of EcN (Fig. 3a, b), yielding the engineered strain EcN-Csg. qRT-PCR assay results revealed that the expression of *csgB* and *csgE* in EcN-Csg were higher than that of EcN, whereas *csgD* was undetectable in EcN-Csg (Fig. 3c). By performing the Congo red-binding assay, our results indicated that the curli fiber formation was enhanced in EcN-Csg as compared to EcN (Fig. 3d). The intestinal adhesion ability of EcN and EcN-Csg were evaluated by incubating these two microbes with MC38 murine colon cells. As shown in Fig. 3e, the intestinal epithelial cells adhesion ability of EcN-Csg was about 2-fold higher than that of EcN.

**Fig. 3.**
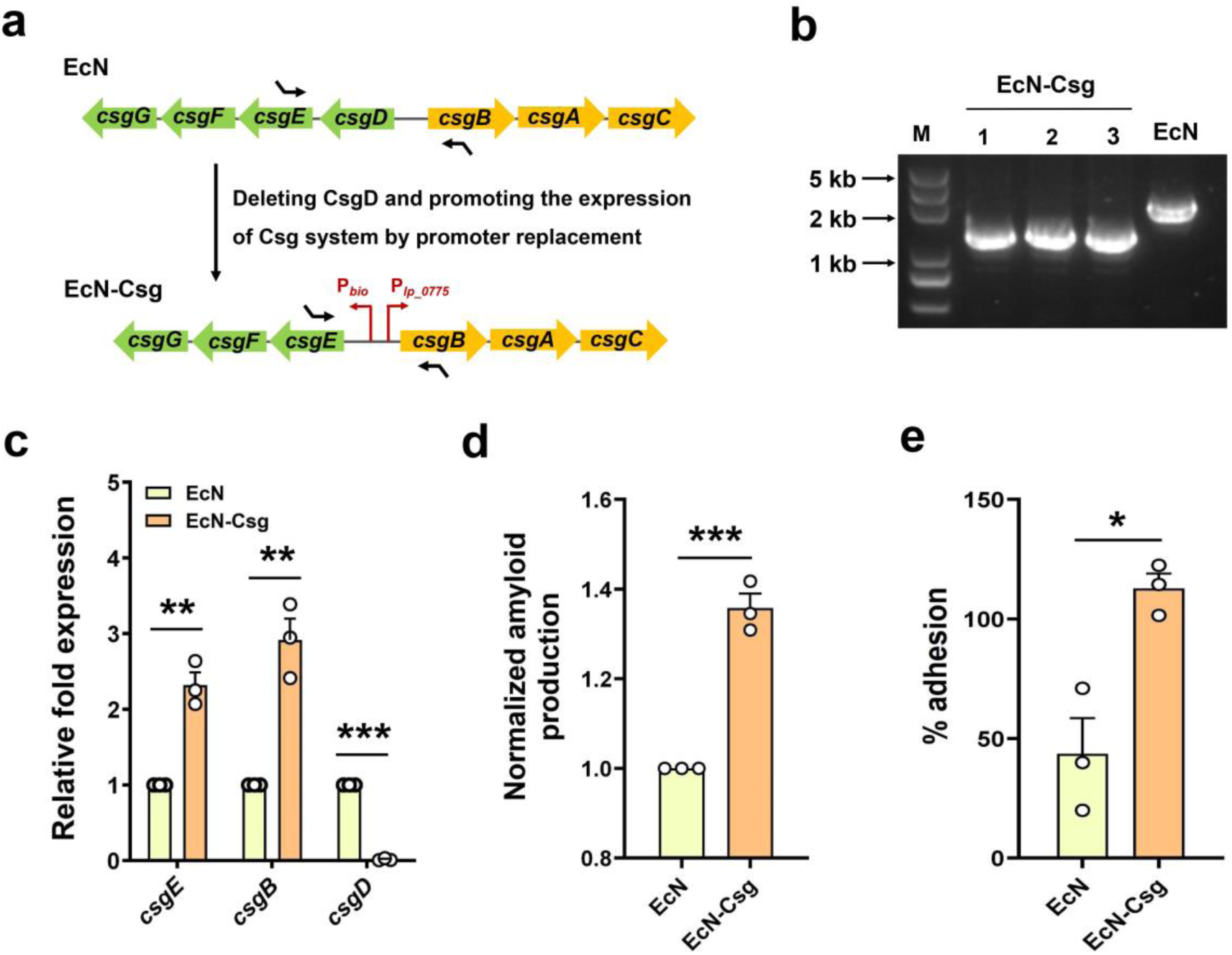
Improving the intestinal colonization of EcN by increasing curli fibers formation. **a** Schematic illustration for the construction of EcN-Csg. **b** PCR verification for EcN-Csg. **c** qRT-PCR analysis for the expression of *csgE*, *csgB*, and *csgD* in EcN and EcN-Csg. **d** Normalized amyloid production of EcN and EcN-Csg. **e** Comparison of the adhesion ability of EcN and EcN-Csg to MC38 cells. The data are presented as mean ± SEM. The statistical significance between two groups were analyzed using *t*-test (\**P* < 0.05, \*\**P* < 0.01, \*\*\**P* < 0.001).

All these results indicated that the intestinal colonization ability of EcN could be improved by strengthening its curli amyloid fiber formation.

### Construction of a synthetic EcN strain with high intestinal colonization ability

Based on the above-mentioned results, we constructed two synthetic EcN strains by integrating the gastric acid and bile salts tolerance-associated element (P*_bio_*-PatE-MdtM) into the *lacZ* gene locus of EcN and EcN-Csg, yielding the engineered strain EcN-PM and Syn3 (Fig. 4a and Supplementary Fig. S3). The *csg* promoter (P*_csg_*) and *lacZ*-targeting sgRNA used in the CRISPR-Cas9 editing technology were listed in Fig. 4a. qRT-PCR analysis revealed that the expression of *patE*, *mdtM*, *csgB*, and *csgE* in Syn3 were higher than that of EcN, whereas the expression of *csgD* was not detected (Fig. 4b). The growth assay results indicated that EcN, EcN-PM, EcN-Csg, and Syn3 showed similar growth curve (Fig. 4c). Scanning electron microscopy (SEM) and Congo red-binding assay results revealed that EcN-Csg and Syn3 produced more curli fibers than that of EcN and EcN-PM (Fig. 4d, e). Besides, the intestinal epithelial cells adhesion ability of EcN-Csg and Syn3 were higher than that of EcN and EcN-PM (Fig. 4f).

**Fig. 4.**
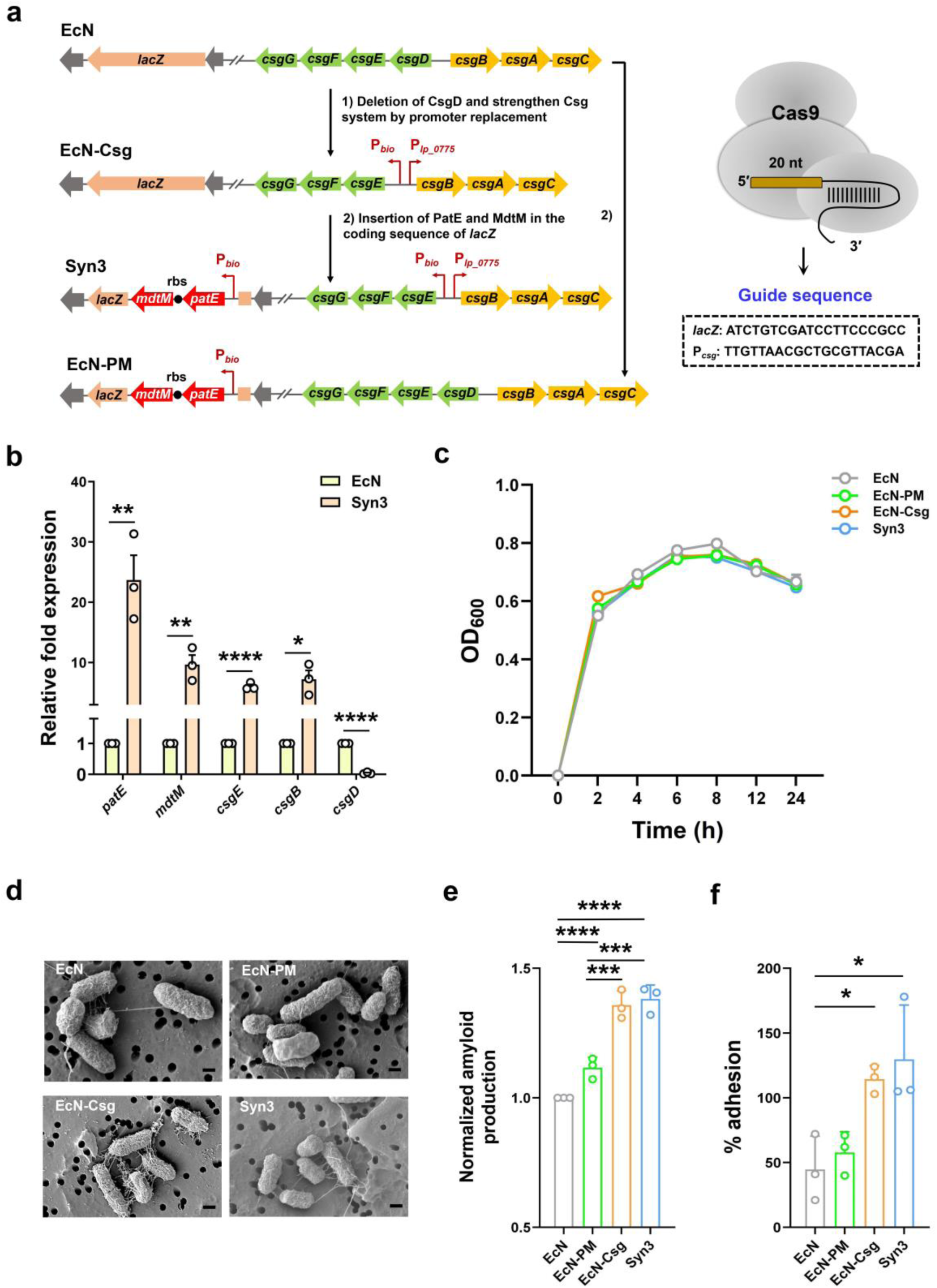
Construction of an engineered EcN strain with high intestinal colonization ability. **a** Schematic illustration for the construction of Syn3 and EcN-PM. **b** qRT-PCR analysis for the expression of *patE*, *mdtM*, *csgE*, *csgB*, and *csgD* in EcN and Syn3. **c** Growth curve of EcN, EcN-PM, EcN-Csg, and Syn3. **d** Electron microscopy-based observation of curli fibers in EcN, EcN-PM, EcN-Csg, and Syn3 (scale bar = 1 μm). **e** Normalized amyloid production of EcN, EcN-PM, EcN-Csg, and Syn3.**f** Comparison of the adhesion ability of EcN, EcN-PM, EcN-Csg, and Syn3 to MC38 cells. The data are presented as mean ± SEM. The statistical significance between two groups were analyzed using *t*-test. The statistical significance between different groups (more than 3 groups) was calculated by using one-way analysis of variance (ANOVA) (\**P* < 0.05, \*\**P* < 0.01, \*\*\**P* < 0.001, \*\*\*\**P* < 0.001).

Next, we tried to determine the intestinal colonization ability of EcN, EcN-PM, EcN-Csg, and Syn3 *in vivo*. To facilitate the visualization and counting of these microbes, the sfGFP-containing plasmid pTD103luxI-sfGFP was introduced into EcN, EcN-PM, EcN-Csg, and Syn3 by electroporation [26]. Then, the C57BL/6J mice were orally administered with PBS and four sfGFP plasmid-containing strains (EcN-fluo, EcN-PM-fluo, EcN-Csg-fluo, and Syn3-fluo) (Fig. 5a). Four hours later, the intact digestive tracts were collected, and the distribution of EcN and EcN-derived strains along the gastrointestinal tract were analyzed by using the *in vivo* imaging system (IVIS). As shown in Fig. 5b, the intestinal colonization ability of Syn3-fluo was higher than the other three strains.

**Fig. 5.**
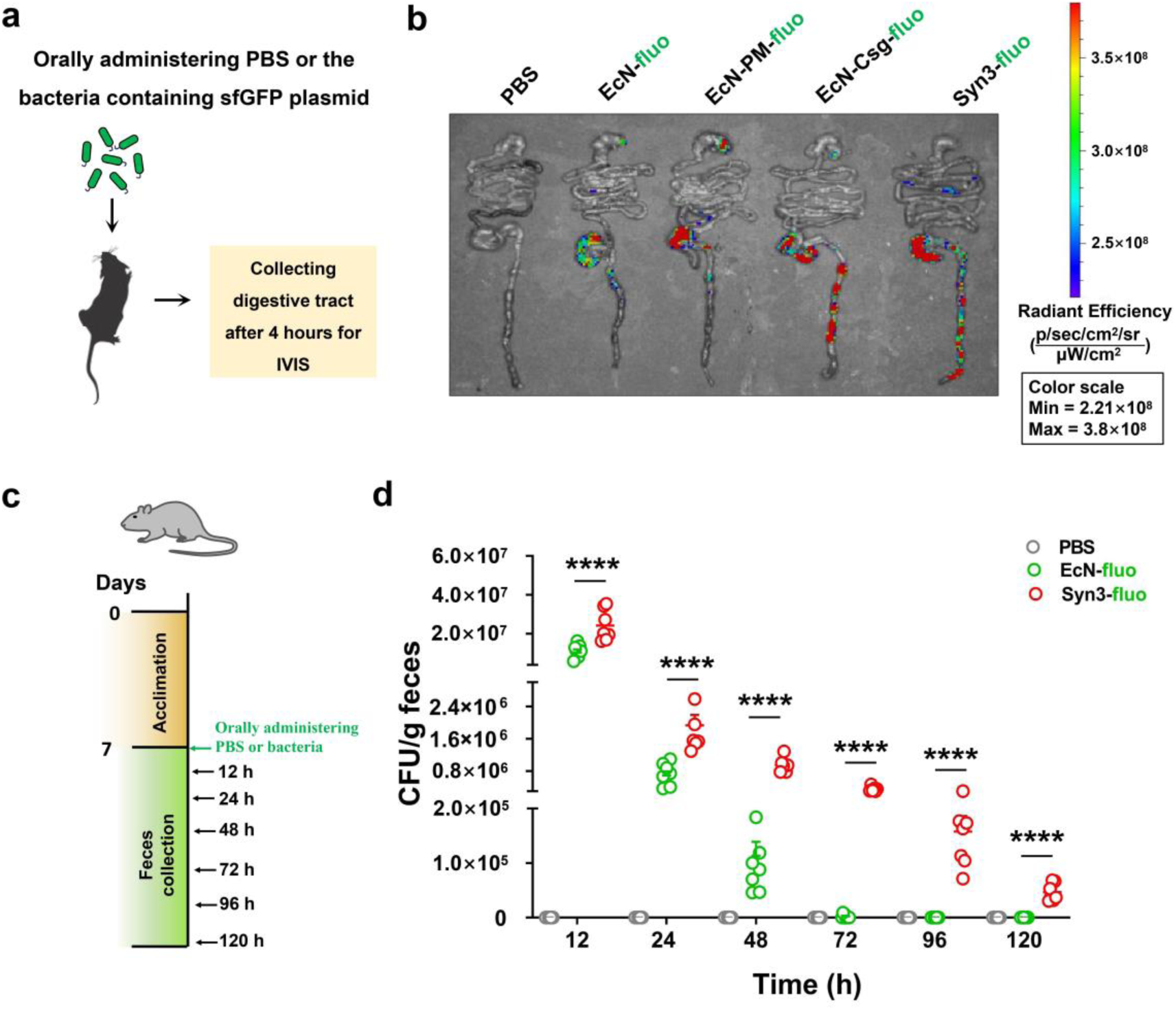
*In vivo* verification of the intestinal colonization ability of Syn3. **a** Schematic illustration of the mouse experiment for the observation of EcN-fluo, EcN-PM-fluo, EcN-Csg-fluo, and Syn3-fluo along the digestive tracts. **b** Bioluminescence images of EcN-fluo, EcN-PM-fluo, EcN-Csg-fluo, and Syn3-fluo in the GI tract of mice. **c** Schematic illustration of the mouse experiment for analyzing the intestinal colonization ability of EcN-fluo and Syn3-fluo. **d** Comparison of the intestinal colonization ability between EcN-fluo and Syn3-fluo. The data are presented as mean ± SEM. The statistical significance between different groups (more than 3 groups) was calculated by using one-way analysis of variance (ANOVA) (\*\*\*\**P* < 0.001).

To further assess the intestinal colonization capacity of Syn3, C57BL/6J mice were orally administered with EcN-fluo and Syn3-fluo at a dose of 1×10^10^ CFUs per mouse (Fig. 5c). The feces were collected at 12, 24, 48, 72, 96, and 120 h post-gavage, dissolved in PBS buffer, and vortexed to acquire the homogenized fecal suspension. After serial dilution, the suspension was spread onto kanamycin-containing LB agar plates and incubated at 37°C for 12 h. As shown in Fig. 5d, Syn3-fluo exhibited higher intestinal colonization ability than EcN-fluo within five days post-gavage.

Both the *in vitro* and *in vivo* results indicated that the intestinal colonization ability of Syn3 was enhanced and superior to EcN.

### Syn3 exhibits higher anti-colitis efficacy than EcN

As reported, EcN administration can alleviate the disease phenotype of colitis by adjusting the disordered gut microbiome and restoring the immune function of the host [27]. Here, EcN and Syn3 were orally administered to the DSS-induced colitis mice to compare their anti-colitis therapeutic efficacy. Specifically, the mice were divided into four groups: (1) NC, the mice provided with normal drinking water and orally administered with PBS buffer containing 20% glycerol; (2) PBS, the mice provided with 3.5% DSS water and orally administered with PBS buffer containing 20% glycerol; (3) EcN, the mice provided with 3.5% DSS water and orally administered with 1 × 10^10^ CFU viable cells of EcN; (4) Syn3, the mice provided with 3.5% DSS water and orally administered with 1 × 10^10^ CFU viable cells of Syn3 (Fig. 6a).

**Fig. 6.**
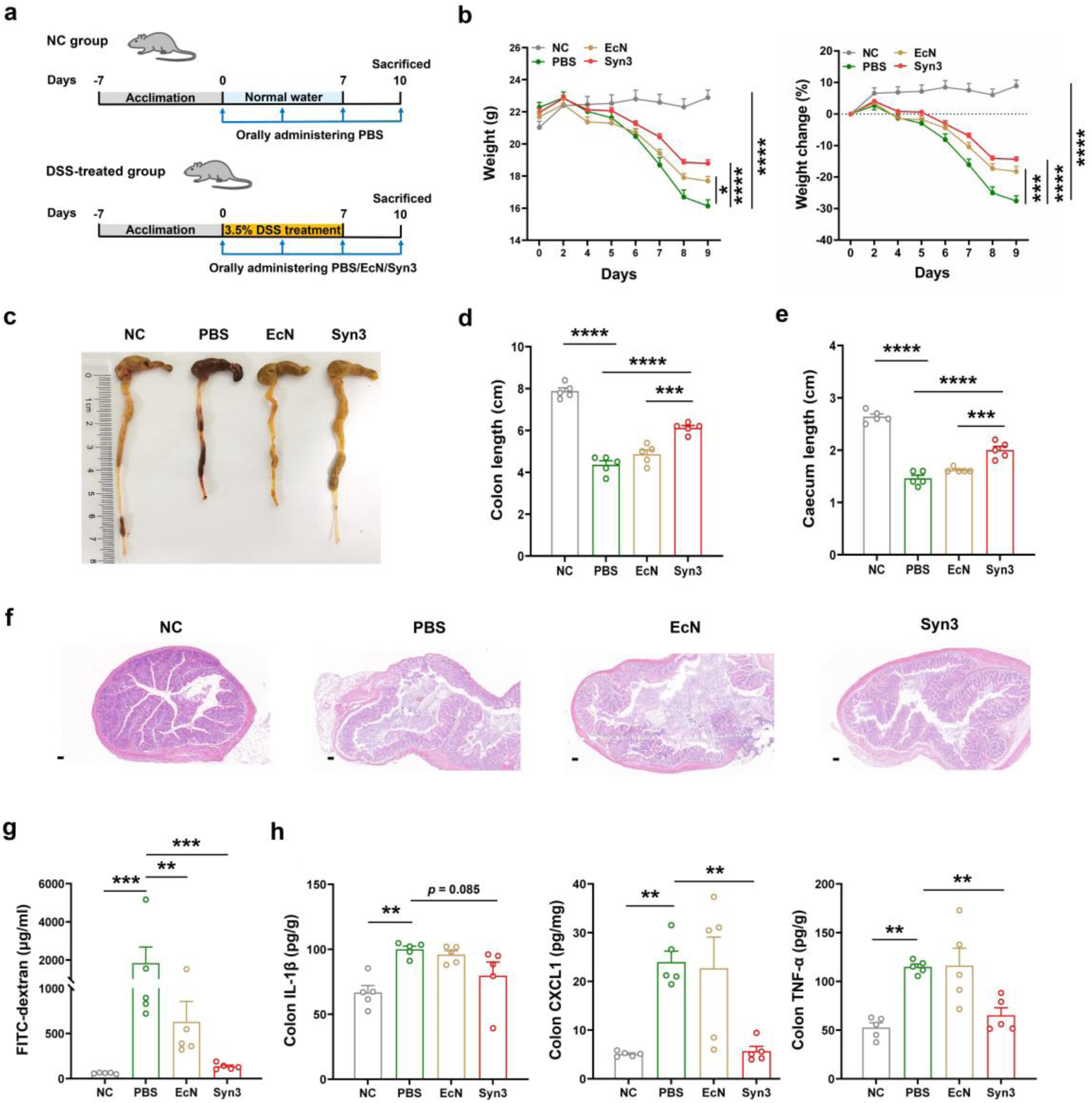
Therapeutic efficacy of Syn3 against DSS-induced colitis. **a** Schematic illustration of the animal experiment. **b** Body weight and weight change of the mice. **c** The photographs of cecum and colon tissues the collected from the mice in different groups at day 9. The length of colon (**d**) and cecum (**e**) in different groups. **f** HE staining analysis of colon tissues that collected from the mice in different groups at day 9. Scale bar: 100 μm. **g** The serum concentration of FITC-dextran in different groups. **h** The concentration of IL-1β, CXCL1, and TNF-α in the colon tissues of different groups. Data are presented as mean ± SEM. The statistical significance between different groups was calculated by using one-way analysis of variance (ANOVA) (\**P* < 0.05, \*\**P* < 0.01, \*\*\**P* < 0.001, \*\*\*\**P* < 0.001).

By evaluating the disease severity of colitis in different groups, we found that the mice of Syn3 group exhibited less weight loss and longer cecum and colon length than that of the PBS and EcN groups (Fig. 6b-e). Hematoxylin-eosin (HE) staining results indicated that the colon sections of Syn3 group showed more intact epithelial cells and regular crypt structure than the PBS and EcN groups (Fig. 6f). To determine the impact of Syn3 administration on the colonic permeability of colitis mice, we performed the fluorescein isothiocyanate (FITC)-dextran perfusion test and found that the intestinal barrier integrity of Syn3 group was superior than that of PBS and EcN groups (Fig. 6g). As for pro-inflammatory cytokines, the concentration of IL-1β, CXCL1, and TNF-α in the colon tissue were increased in the PBS and EcN groups, whereas decreased in the Syn3 group (Fig. 6h).

All these results revealed that Syn3 administration exerts superior colitis-protective effects than EcN as evidenced by improving intestinal barrier integrity, reducing inflammatory responses, and preserving colonic morphology.

### Oral administration of Syn3 restored the disordered gut microbiome of colitis mice

Considering that gut microbiota is closely associated with the progression of ulcerative colitis (UC), and microbial intervention has been proved to be an effective strategy for treating colitis [28], we performed the 16S rRNA amplicon sequencing to verify whether Syn3 administration could restore the disordered cecal and colonic microbiome of colitis mice. The number of 16S rRNA gene amplicon sequencing reads of each sample was rarefied to 21,599, yielding an average Good’s coverage of over 99.68 % and 99.83% in the cecum and colon samples, respectively (Supplementary Fig. S4a, b). Venn diagram analysis revealed that 131 amplicon sequence variants (ASVs) were shared by the cecal microbiome of all four groups, while 939, 468, 610, and 300 ASVs were uniquely present in the NC, PBS, EcN, and Syn3 groups (Fig. 7a). Differently, 92 ASVs were shared by the colonic microbiome of four mice groups, while 805, 297, 123, and 224 ASVs were uniquely observed in the NC, PBS, EcN, and Syn3 groups (Fig. 7b). Consistent with the previously reported findings [29], the richness (Ace) and diversity (Shannon) of cecal and colonic microbiome decreased significantly in the colitis mice groups (PBS, EcN, and Syn3) as compared to the NC group (Fig. 7c, d). As for the gut microbial dysbiosis index, although it was increased in the cecal and colonic samples of three colitis mice groups (PBS, EcN, and Syn3), Syn3 administration could reduce its level to some extent, especially for the cecal samples (Supplementary Fig. S4c, d). Principal coordinate analysis (PCoA) of weighted UniFrac distances showed that the cecal and colonic microbiome composition of the PBS, EcN, and Syn3 groups were different from that of the NC group (Fig. 7e, f). In order to examine the cecal and colonic microbial differences between the four mice groups, the top 50 genera were selected and used to perform the heatmap analysis (Fig. 7g, h). Linear discriminant analysis effect size (LEfSe) analysis (LDA score >2.0) indicated that the cecal microbes that belonging to *norank_f_Muribaculaceae* and *NK4A214_group* were enriched in the NC group, while the colonic microbes that increased in the NC group belonged to *Lactobacillus*, *Parvibacter*, and *norank_o_Clostridia_UCG-014*; the cecal and colonic microbes belonging to *Escherichia-Shigella* and *Parabacteroides* were enriched in the PBS group; the cecal microbes belonging to *Akkermasia*, *Butyricicoccus*, *Acetatlfactor*, and *Romboutsia* were abundant in the EcN group, while the colonic microbes enriched in this group belonged to *Enterococcus* and *Romboutsia*; the cecal microbes belonging to *Parvibacter*, *Turicibacter* and *Family_XIII_ad3011_group* were abundant in the Syn3 group, whereas the colonic microbes of *Tricibacter*, *norank_f_Muribaculaceae*, and *Streptococcus* enriched in this group (Supplementary Fig. S4e, f).

**Fig. 7.**
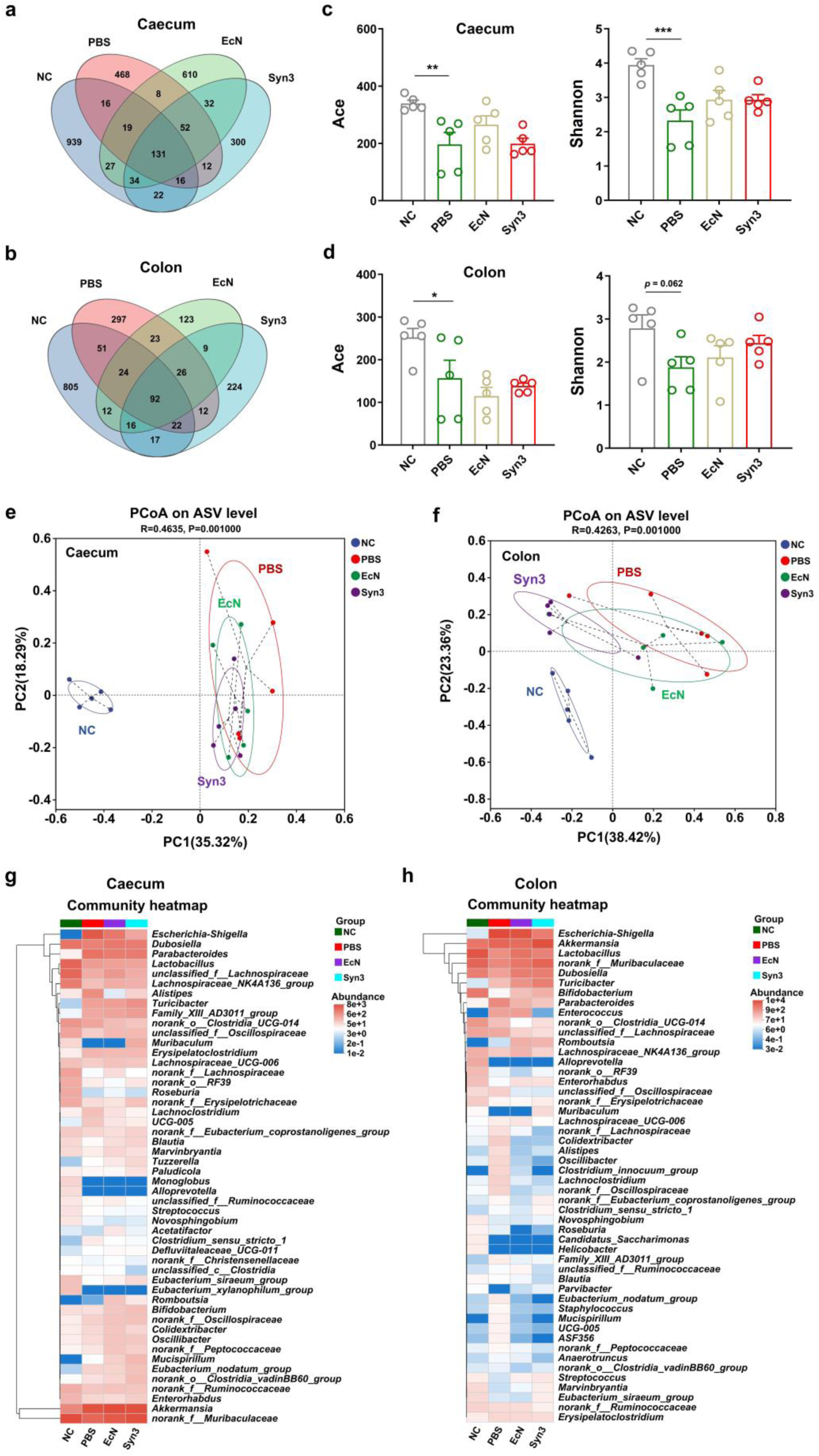
Syn3 administration affects the cecal and colonic microbiome of colitis mice. Venn diagram showing the number of ASVs in the cecum (**a**) and colon (**b**) microbiome of four mice groups. The richness (Ace) and diversity (Shannon) of cecal (**c**) and colonic (**d**) microbiome in four mice groups. PCoA of weighted UniFrac distances based on the ASVs showing the cecal (**e**) and colonic (**f**) microbiome differences among the four mice groups. Community heatmap analysis for the comparison of top 50 bacterial genera in the cecal (**g**) and colonic (**h**) microbiome among the four mice groups. Data are presented as mean ± SEM. The statistical significance between different groups was calculated by using one-way analysis of variance (ANOVA) (\**P* < 0.05, \*\**P* < 0.01, \*\*\**P* < 0.001).

Next, we tried to analyze the relative and absolute abundances of cecal and colonic microbiome at the genus level. To fulfill this goal, we quantified the fecal bacterial load using a 16S rRNA quantitative PCR assay, and found that the cecal and colonic bacterial load showed no significant differences among the four mice groups (Supplementary Fig. S5a, b). Then, the inferred absolute abundance of specific genus was calculated by multiplying the fecal bacterial load and its relative abundance. Syn3 administration could decrease the relative and inferred absolute abundance of *Escherichia-Shigella* in the cecal and colonic samples (Fig. 8a, b). As for the genus *norank_f_Muribaculaceae*, its relative abundance was increased in the Syn3 group as compared to the PBS group in the cecal and colonic samples. Differently, the inferred absolute abundance of cecal and colonic microbes belonging to *norank_f_Muribaculaceae* showed no significant differences between the four mice groups. Apart from the genera mentioned above, the relative abundance of *Streptococcus* was increased in the EcN and Syn3 groups as compared to the PBS group in the colonic sample; the relative abundance of colonic microbes belonging to *UCG-005* was decreased in the EcN and Syn3 groups as compared to the PBS group (Fig. 8a, b).

**Fig. 8.**
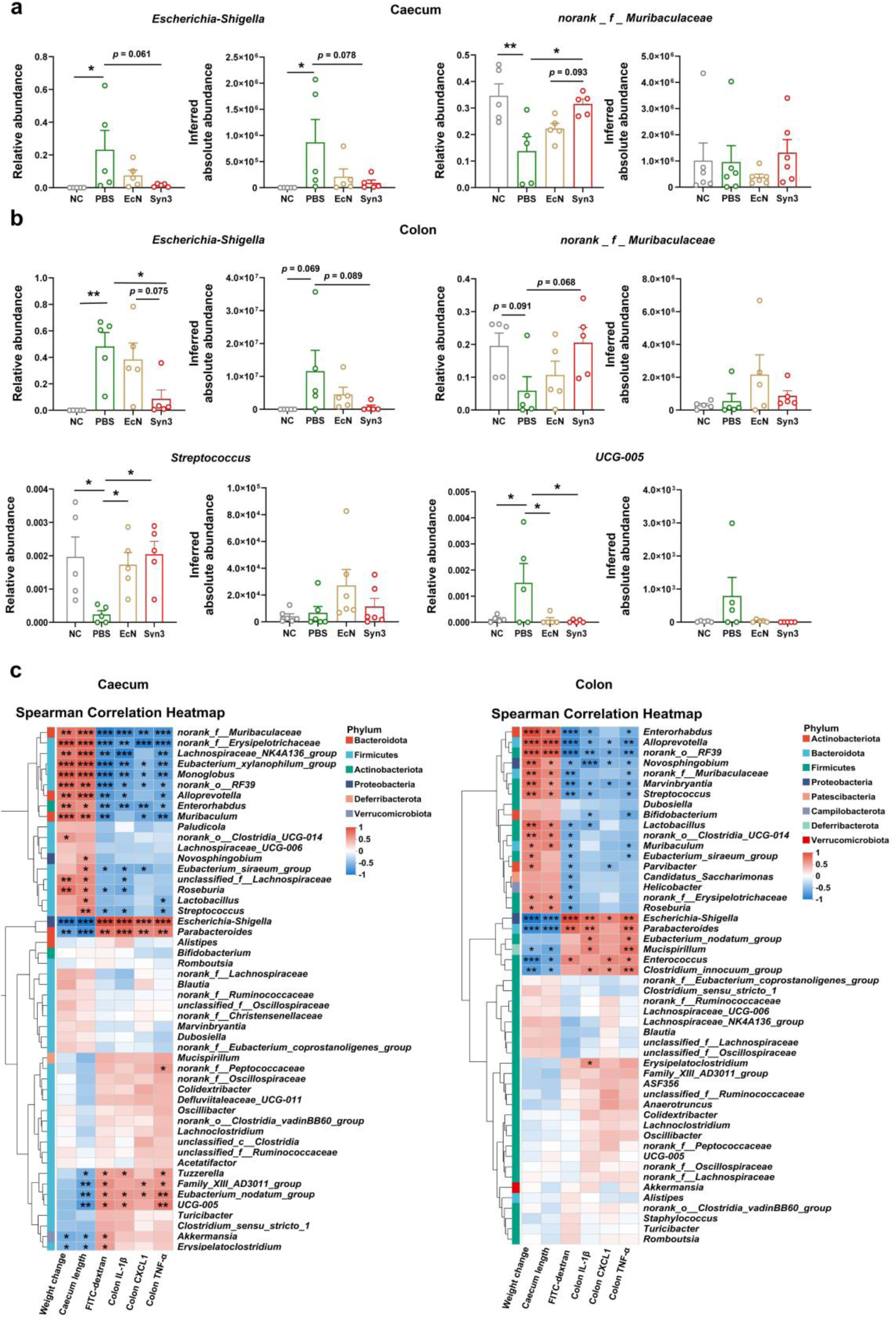
Syn3 administration affected cecal and colonic microbial genera and their correlations with colitis-associated phenotypes. Relative and inferred absolute abundances of cecal (**a**) and colonic (**b**) microbiome at genus level. The correlations between the cecal (**c**) and colonic (**d**) microbes and colitis-associated phenotypes. Data are presented as mean ± SEM. The statistical significance between different groups was calculated by using one-way analysis of variance (ANOVA) (\**P* < 0.05, \*\**P* < 0.01).

We performed the Spearman correlation analysis to predict the interactions between the genera abundance and colitis-associated phenotypes. As shown in Fig. 8c, the cecal and colonic microbes from *Escherichia-Shigella*, and *Parabacteroides* were responsible for the inflammatory responses and intestinal damage of colitis mice; the cecal and colonic microbes from *norank_f_Muribaculaceae*, *Muribaculum*, *norank_o_RF39*, *Enterorhabdus*, and *Alloprevotella* were positively correlated to the beneficial indicators, including reduced inflammation, preservation of intestinal length, integrity of intestinal barrier, and attenuation of weight loss. These results suggested that changes in the relative abundance of the aforementioned microbes in the cecal and colonic microbiome may influence colitis progression.

In summary, our results indicated that Syn3 administration can restore the disturbed cecal and colonic microbiome by decreasing *Escherichia-Shigella* and increasing *norank_f_Muribaculaceae*, which might contribute to the alleviation of colitis phenotypes.

### *In vivo* safety evaluation of Syn3

To evaluate the biosafety of Syn3 for potential clinical application in colitis therapy, we administered it by oral gavage to C57BL/6J mice every three days over a 30-day period. (Supplementary Fig. S6a). The mice that treated with PBS solution containing 20% glycerol were used as control (PBS gavage). The healthy state of mice in the Syn3-treated group (Syn3 gavage) were examined by checking the variations of various metabolic and physiological indicators. Both the weight and weight change of the Syn3-treated mice were comparable to that of the PBS-treated mice (Supplementary Fig. S6b). As for the food and water intake, there were no significant differences between the two mice groups (Supplementary Fig. S6c). We examined the mental status and behavioral activity of mice by measuring the serum dopamine level and performing the tail-hanging and pole-climbing tests. Our results indicated that Syn3 administration had no significant effect on neurobehavioral function or motor activ5ity (Supplementary Fig. S6d-f). Besides, we performed the routine blood and biochemical analyses to compare the physiological parameters between the two mice groups. As shown in Supplementary Fig. S6g, the concentration of white blood cells, neutrophils, lymphocytes, and eosinophils were comparable between the two mice groups. Differently, the concentration of monocytes was increased slightly in the Syn3-treated mice. As for the anaemia-related indicators, the hemoglobin (HGB) and mean corpuscular hemoglobin concentration (MCHC) of the Syn3-treated mice were comparable to that of the control mice (Supplementary Fig. S6h). We compared the liver function of the PBS and Syn3-treated mice by assaying the serum concentration of glutamic-oxaloacetic transaminase (GOT), and found that it showed no significant difference between the two mice groups (Supplementary Fig. S6i). Apart from the parameters mentioned above, our results revealed that the coefficients of main organ (colon length, cecum length, and spleen/liver/kidney/brain/thymus/testis) of the Syn3-treated mice were comparable to that of the PBS-treated mice (Supplementary Fig. S6j, k). All these results indicated that Syn3 administration exhibited no side effects on the health of C57BL/6J mice.

## Discussion

The harsh GI environment, particularly the presence of gastric acid and bile acids, can restrict the intestinal colonization of EcN, thus hindering the application of EcN-derived synthetic probiotics for disease treatment [8]. In this study, we engineered EcN to improve its intestinal colonization ability by enhancing the expression of gastric acid and bile acids tolerance-associated genes and strengthening the formation of curli amyloid fibers. Oral administration of this engineered strain to the colitis mice alleviated most of the disease phenotypes, which might be achieved by restoring the disordered gut microbiome of colitis mice. Based on the high intestinal colonization ability and colitis-protective properties of this engineered strain, our study provided an excellent and stable chassis for the construction of EcN-derived synthetic probiotics for disease treatment.

To improve the bile acids tolerance of EcN, we overexpressed two bile acids exportation-associated multidrug resistance transporters MdtM and DinF in EcN. Our results indicated that MdtM could transport cholate and deoxycholate efficiently, while DinF was only responsible for the transportation of deoxycholate (Fig. 2d), indicating that DinF might be a deoxycholate-specific transporter, which need to be further verified by applying more bile acids. This result further highlights the diverse bile acid transport capabilities of EcN.

It had been reported that overexpression of heterologous proteins in bacterial systems might impose a substantial metabolic burden on host cells, resulting in significantly reduced growth rates. In this study, utilizing CRISPR-Cas9-based genome editing technology, all the functional elements were integrated into the genome of EcN, yielding a synthetic probiotic with stable inheritance and maintenance of functional genes while preserving normal growth ability (Fig. 4a, c). Compared with the strategies that improving the intestinal survival rate of EcN by encapsulating it with protective and adhesive materials [13, 14], our engineered strain avoided the time-consuming procedure of encapsulation and exhibited prolonged intestinal duration time and high anti-colitis therapeutic efficacy (Figs. 5 and 6).

The dysbiosis of gut microbiome was closely associated with the progression of IBD [30]. As reported, EcN administration could alleviate IBD by inhibiting the growth of pathogenic bacteria such as *Salmonella* [31], *Staphylococcus aureus* [32], and *Shigella* [33], enhancing intestinal tight junction [34, 35], and modulating the intestinal immune function [36].

Consistent with these results, our study revealed that oral administration of Syn3 exhibited higher anti-colitis effect than EcN by sustaining the integrity of the intestinal barrier (Fig. 6f, g) and decreasing the levels of pro-inflammatory factors IL-1β, CXCL1 and TNF-α (Fig. 6h). Besides, Syn3 administration could restore the cecal and colonic microbiome of colitis mice by decreasing the abundance of *Escherichia-Shigella*, while increasing *norank_f_Muribaculaceae* (Fig. 8a, b). As reported, the pathogenic microbes of *Escherichia-Shigella* could exacerbate colitis by disrupting the intestinal barrier, exacerbating inflammation, and triggering immune responses [37, 38], while *norank_f_Muribaculaceae* could alleviate colonic inflammation by producing the anti-inflammatory short chain fatty acids (SCFAs), repairing the intestinal barrier, inhibiting the growth of pathogenic microbes, and modulating the expression of mucin genes [39].

Thus, the Syn3-induced reduction of *Escherichia-Shigella* and increase of *norank_f_Muribaculaceae* might jointly contributed to the alleviation of colitis symptoms. Notably, these microbial shifts were more pronounced in the Syn3 group than in the EcN group. (Fig. 8a, b), which was consistent with the superior anti-colitis effect of Syn3 (Fig. 6). Although our results deciphered the gut microbial alternations following Syn3 administration, however, the variation in gut microbiota was determined by 16S rRNA gene sequencing, which only accurately reflect the microbial variations at genus level. Thus, future studies will employ metagenomic sequencing to identify the specific bacterial species affected by Syn3 administration.

In summary, our study revealed that the probiotic properties of EcN could be improved by enhancing its resistance towards gastric acid and bile salts and its intestinal adhesion ability. By integrating all functional elements into the EcN genome, we have constructed a synthetic EcN strain with high probiotic properties and excellent anti-colitis efficacy, thus providing an excellent and stable chassis for the construction of EcN-derived synthetic probiotics for the treatment of various diseases.

## Materials and Methods

### Strains, plasmids, and growth conditions

The Top 10 were used to clone the genes. *E. coli* Nissle 1917 (EcN) was obtained from Pharma-Zentrale GmbH (Germany), grown in the LB medium or on LB agar under aerobic or anaerobic conditions, and supplemented with 100 μg/ml ampicillin, 50 μg/ml spectinomycin, or 50 μg/ml kanamycin when necessary. The anaerobic growth and fermentation of EcN were performed using an anaerobic chamber (Electrotek AW500TG-D, UK). The strains and plasmids used in this work are listed in Supplementary Table 1 and Supplementary Table 2.

The fragments of P*_adc_* (promoter sequence of CAP0165), P*_lp_0237_*, P*_CAC3456_*, P*_lp_0075_*, P*_sol_* (promoter sequence of CAP0162-0164), and P*_bio_* (promoter sequence of CAC1360-1362) were PCR-amplified by using the genome of *Clostridium acetobutylicum* and *Lactobacillus plantarum* as template. After digestion with *Kpn*I and *Nde*I, the fragments were cloned into the plasmid pET-32a-*lacZ*, yielding the plasmids pET-32a-P*_adc_*-*lacZ*, pET-32a-P*_lp_0237_*-*lacZ*, pET-32a-P*_CAC3456_*-*lacZ*, pET-32a-P*_lp_0775_*-*lacZ*, pET-32a-P*_sol_*-*lacZ*, and pET-32a-P*_bio_*-*lacZ*; The plasmid pET-32a-P*_bio_* was construct by replacing P*_thl_* with P*_bio_* in pET-32a-P*_th_*_l_ [40]; The genes *patE* (CIW80_12240), *dinF* (CIW80_15360) and *mdtM* (CIW80_17355) were PCR-amplified by using the genome of EcN as template. The fragments were digested with *Sal*I and *Nde*I and cloned into the plasmid pET-32a -P*_bio_*, yielding the plasmids pET-32-P*_bio_*-*patE*, pET-32a-P*_bio_*-*dinF*, and pET-32a-P*_bio_*-*mdtM*. For the construction of pET-32a-P*_bio_*-*patE*-*mdtM*, *patE* and *mdtM* were PCR-amplified and ligated together by overlap PCR, yielding the fragments *patE-mdtM*. To be noted, an RBS sequence (TTTAAAGGGAGGATATGAAC) was inserted between *patE* and *mdtM* to improve the translational efficiency of MdtM. Then, the *patE-mdtM* fragment was digested with *Sal*I and *Nde*I and cloned into the plasmid pET-32a-P*_bio_*, yielding pET-32a-P*_bio_*-*patE-mdtM*. The genome-editing of EcN was accomplished by applying the CRISPR-Cas9 system as reported previously [41].

### Animals

The mice were housed under specific pathogen-free conditions in a controlled environment with a 12-hour light–dark cycle, maintained at a temperature of 20°C and a relative humidity of 40-60%. Standard chow diet (P1101F-25, Pulutengbio, China) and water were provided ad libitum. The animal procedures adhered to the guidelines set by the Animal Care Committee of the Shanghai Institute of Biological Sciences, the Institute of Neuroscience, and the Chinese Academy of Sciences Center for Excellence in Brain Science and Intelligence Technology (Approval No. NA-041-2022-R1).

### β-galactosidase assay

The plasmids pET-32a-P*_adc_*-*lacZ*, pET-32a-P*_lp_0237_*-*lacZ*, pET-32a-P*_CAC3456_*-*lacZ*, pET-32a-P*_lp_0775_*-*lacZ*, pET-32a-P*_sol_*-*lacZ*, and pET-32a-P*_bio_*-*lacZ* were electroporated into EcN and grown in LB medium containing 50 μg/ml ampicillin under both aerobic and anaerobic conditions. After incubation at 37°C for 12 h, the cell pellets were harvested by centrifugation (12,000 g, 4°C, 10 min), dissolved in B-PER reagent (Thermo Scientific Pierce®, USA) and vortexed for 1 min to lyse cells. The cell lysate was heat treated at 60°C for 30 min to remove the heat-unstable proteins. After centrifugation at 12,000 g for 30 min, the supernatant was collected and used for β-galactosidase assays as previously reported [40, 42].

### Acid resistance assay

To investigate whether PatE overexpression could improve the acid tolerance of EcN via the AR1-3 systems, we performed the acid tolerance assay as previously reported [18, 43]. For the AR1-mediated acid tolerance assay, EcN-P (the control strain harboring an empty vector) and EcN-PatE (*patE* overexpression strain) were cultivated overnight in LB broth (pH 7.0) at 37°C under constant agitation (200 rpm). The grown cells were inoculated into the same medium and grown for 8 to 10 hours until OD_600_ reach 1.0. Then, the grown cells were diluted (1:10) in the LB medium with the pH reached 2.5 and incubated at 37°C for 2 h under constant agitation (200 rpm). The number of bacterial cells before and after acid challenge were determined by spreading the collected cell cultures onto LB plates. The fold change survival of specific bacteria was calculated by dividing the bacterial numbers after challenge by the bacterial numbers before challenge. For the AR2- and AR3-based acid tolerance assay, EcN-P and EcN-PatE were grown in EG medium [44] (pH 7.0) at 37°C under constant agitation (200 rpm). Then, the grown cells were diluted 1:10 in EG medium (pH 2.5) supplemented with 1.5 mM glutamate (AR2) or 1.5 mM arginine (AR3). After incubation at 37°C for 2 h under constant agitation (200 rpm), the bacterial viability and fold change survival was determined as described above.

### Bile salts resistance assay

To assess the bile salts resistance of EcN-P (the control strain harboring an empty vector), EcN-MdtM (*mdtM* overexpression strain), and EcN-DinF (*dinF* overexpression strain), we performed the bile salts resistance assay as previously reported [21]. Specifically, the bacteria cells were grown in LB liquid medium containing various concentrations of sodium salts of cholate (8, 12, 16, 24, 32 and 40 mg/ml) and deoxycholate (4, 8, 12, 24, 32, 48 and 64 mg/ml) (pH = 7.2). IC_50_, the concentration of bile salts that inhibit the growth of bacterial cells by 50%, was calculated by using the GraphPad Prism software and used to quantify the bile salts resistance of EcN-P, EcN-MdtM, and EcN-DinF.

### Gastrointestinal-tract mimicking assay

The gastrointestinal-tract survival analysis was performed as previously reported [23]. EcN-PatE-MdtM (*patE* and *mdtM* overexpression strain) and EcN-P (the control strain harboring an empty vector) were grown in LB medium at 37℃. The bacterial cells harvested at the logarithmic phase were washed three times with pre-warmed (37°C) PBS solution, resuspended in pre-warmed (37°C) PBS buffer containing gastric juice (53 mM NaCl, 15 mM KCl, 5 mM Na_2_CO_3_, 1 mM CaCl_2_, 0.1 mg/ml lipase (Sigma, ^L^3126,USA), and 1.2 mg/ml pepsin (Sigma,P6887,USA), pH=2.4), and incubated on a slow-moving rotary shaker (10 rpm) at 37°C for 60 minutes. Then, the pH of cultures was neutralized to 6.5 by adding NaHCO_3_, followed by the addition of pancreatic juice (85 mM NaCl, 5 mM KH_2_PO_4_, 2 mM Na_2_HPO_4_ and 10 mM NaHCO_3_) containing pancreatin (30 mg/ml) (Sigma, P1750,USA) and bile acids (3.0 mM sodium glycocholate hydrate, 1.3 mM sodium glycodeoxycholate, 2.4 mM sodium glycochenodeoxycholate, 1.0 mM taurocholic acid sodium salt hydrate, 0.4 mM sodium taurodeoxycholate hydrate, and 1.0 mM sodium taurochenodeoxycholate). The mixture was incubated for another 30 minutes. The bacterial suspensions that collected at 0, 30, 60, and 90 min were serial diluted and spread on the LB plate to calculate the CFUs of EcN-PatE-MdtM and EcN-P. Then, the relative survival of EcN-PatE-MdtM and EcN-P at 30, 60 and 90 minutes were calculated as the following equation: CFUs (30, 60 and 90 minutes)/ CFUs (0 minutes).

### Quantitative real-time RT-PCR (qRT-PCR)

To compare the expression of acid tolerance-associated genes between EcN-P (the control strain harboring an empty vector) and EcN-PatE (patE overexpression strain), these two bacteria were inoculated into LB medium containing 50 μg/ml ampicillin. After incubation at 37°C for 9 h, the grown cells were collected. To compare the expression of curli fiber formation-associated genes between EcN and EcN-Csg, these two bacteria were inoculated into LB medium. After incubation at 37°C for 9 h, the grown cells of these two bacteria were collected. To examine the expression level of the functional genes that integrated into the genome of Syn3, EcN and Syn3 were inoculated into LB medium. After incubation at 37°C for 9 h, the grown cells of these two bacteria were collected.

The RNA of collected bacteria was isolated by TRIZOL extraction as described previously [45]. Then, RNA was reverse transcribed to cDNA using the PrimeScript RT Reagent Kit (TaKaRa, cat. #RR047A, China). qRT-PCR was carried out by using the ChemQ SYBR color qPCR Master Mix (Vazyme, Q421-02-AA, China) with the following conditions: 95°C for 2 min, followed by 40 cycles of 95°C for 15 s, 55°C for 20 s, and 72°C for 20 s. The relative abundance of transcripts was calculated by normalizing to that of 16S rRNA. The primers that were used for qRT-PCR analysis are listed in Supplementary Table 3.

### Cell adhesion assay

Murine Carcinoma-38 (MC38) cells were obtained from Kerafast (ENH204-FP, Boston) and cultured in Dullbecco’s Modified Eagle’s Medium (DMEM) (Gibico, 11,965,092, USA) at 37°C in a 5% CO_2_-containing humidified atmosphere. Microbes were inoculated into LB medium and grown at 37°C anaerobically until the OD_600_ reached 0.6. Then, the grown bacterial cells were collected by centrifugation, washed with PBS solution for three times, and dissolved in the PBS solution. Then, 1 × 10^7^ CFU viable cells of each microbe was incubated with MC38 cells (cell amount=1×10^5^) at a ratio of 100:1 at 37°C in a 5% CO_2_-containing humidified atmosphere. After incubation for 3 h, the supernatants were discarded. The cells were washed with PBS solution three times and treated with 0.2% Triton-X100. The cell lysates were diluted, spread onto an LB solid medium, and incubated overnight at 37°C. The CFUs of microbes were counted. The cell adhesion rate of bacteria was calculated using the following formula: (the number of microbes in the bacteria-treated group minus the number of bacterial cells in the PBS-treated group) ×dilution fold/number of MC38 cells.

### Electron microscopy-based observation of curli fibers

EcN, EcN-PM, EcN-Csg, and Syn3 were inoculated into 5 ml LB liquid medium and grown at 37°C until OD_600_ reached 1.0. Then, 2 ml of the cultures were collected and fixed on nucleopore track-etched membranes (Whatman, 110611, UK) by using a microsyringe filter (Merck, XX3002500, USA). The bacteria-containing membranes were immersed in the fixative solution (2% glutaraldehyde and 2% paraformaldehyde in 0.1 M sodium cacodylate buffer) for 24 h at 4℃. The membranes were gently washed with PBS buffer, dehydrated with ethanol, dried by using the critical point dryer (Leica, EM CPD300, Germany), and coated with 80:20 Pt: Pd (10 nm thick). Finally, the field emission scanning electron microscope (Zeiss, FE-SEM ZeissGemin300, Germany) was used to image the curli fibers of different bacteria.

### *In vivo* verification of the intestinal colonization ability

The four-week-old male C57BL/6J mice were purchased from Charles River Laboratories China, maintained under pathogen-free conditions on a 12 h light-dark cycle at a temperature of 24°C and 40-60% humidity, and kept free access to food and water. Before the experiment, all the mice experienced a one-week acclimation period. To facilitate the visualization of bacteria, the sfGFP-containing plasmid pTD103luxI-sfGFP was introduced into EcN, EcN-PM, EcN-Csg, and Syn3 by electroporation, yielding the recombinant strain EcN-fluo, EcN-PM-fluo, EcN-Csg-fluo, and Syn3-fluo. Then, these bacteria were orally administered to the C57BL/6J mice that fasted for 4 hours before the experiment (1×10^10^ CFUs per mouse). Four hours later, the intact digestive tracts were collected, and the distribution of EcN and EcN-derived strains along the gastrointestinal tract was observed by using PerkinElmer IVIS Spectrum (Excitation light at 460-520nm). The living image software (PerkinElmer, version 4.5) was used for image analysis.

To verify the intestinal colonization ability of EcN-fluo and Syn3-fluo, these two bacteria were orally administered to the C57BL/6J mice. In this experiment, the mice were divided into three groups: (1) PBS (*n* = 8), the mice that intragastric administered with PBS buffer containing 20% glycerol; (2) EcN-fluo (*n* = 8), the mice that intragastric administered with EcN-fluo (1.0×10^10^ CFUs per mouse); (3) Syn3-fluo (*n* = 7), the mice that intragastric administered with Syn3-sfGFP (1.0×10^10^ CFUs per mouse). The feces were collected at 12, 24, 48, 72, 96, and 120 h post-gavage. The collected feces were dissolved in PBS buffer and vortexed for 5 min to get the homogenized solution. Then, the samples were diluted with PBS buffer and spread on the LB agar plates containing 50 μg/ml kanamycin. The green colonies were counted and used to evaluate the intestinal colonization ability of EcN-fluo and Syn3-fluo at different time points.

### Congo Red-binding assay-based quantification of amyloid production

EcN, EcN-PM, EcN-Csg, and Syn3 were inoculated into 1 ml LB liquid medium and grown at 37°C until OD_600_ reached 1.0. After centrifugation, the collected cells were suspended in 0.025 M Congo Red (CR) solution and incubated for 10 min. Then, the supernatants were collected by centrifugation (14,000 rpm, 10 min). The absorbance of the supernatant at 490 nm was measured by using the Variokan LUX multifunctional enzyme labeler (Thermo Fisher Scientific, USA). Normalized curli fiber production was calculated by subtracting the measured absorbance value from that measured for 0.025 mM Congo Red in PBS and normalized by the OD_600_ of the original bacterial culture.

### Histological analysis of colon tissue

The distal colons were collected and fixed in 4% paraformaldehyde for 24 h. Then, the colon tissue was embedded in paraffin, sectioned (3 μm), and stained with a hematoxylin and eosin solution. Finally, the sections were imaged using a Panoramic MIDI slide scanner (3D HISTECH, Pannoramic SCAN II, Hungary).

### Construction of DSS-induced colitis mice for disease treatment

Eight-week-old male C57BL/6J mice were purchased from Charles River Laboratories. After one week acclimation period, the mice were divided into four groups (n=9-12): (1) NC, the mice that received normal water drinking and intragastric administration of PBS buffer containing 20% glycerol; (2) PBS, the mice that received 3.5% dextran sulfate sodium salt (DSS, Meilunbio, MB5535,China) and intragastric administration of PBS buffer containing 20% glycerol; (3) EcN, the mice that received 3.5% DSS and intragastric administration of EcN (1.0×10^10^ CFUs per mouse); (4) Syn3, the mice that received 3.5% DSS and intragastric administration of Syn3 (1.0×10^10^ CFUs per mouse). For the DSS-induced colitis mice, they were fed with drinking water containing 3.5% DSS for 7 days, and then replaced with normal water during the following days. The oral administration of bacterial cells was performed every three days.

During the experiment, the body weight of mouse was measured every day. After 9 days treatment, the mice were executed to measure the cecum and colon length and collect different tissues.

### Assay of epithelial integrity

FITC-dextran assay was used to evaluate intestinal integrity and performed as previously reported [46]. In brief, the mice that deprived of food and water for 4 h before the experiment were oral administered with FITC-dextran (0.5mg/g) (Chondrex, 4013, USA). After 3 h post-gavage, the serum was collected to detect the FITC fluorescence signal with microplate reader (Excitation: 490 nm, Emission: 520 nm).

### ELISA-based measurement of pro-inflammatory factor in colon tissue

To detect the level of pro-inflammatory cytokines, the colon tissues of mice were collected, weighed, and homogenized by using the cold PBS buffer. The concentrations of IL-1β, TNF-α, and CXCL1 of colon tissue were determined by ELISA kits (IL-1β, Absin, abs520001-96T,China; TNF-α, Absin, abs552204-96T,China; CXCL1, Absin, abs520017,China) following the manufacturer’s instructions.

### 16S rRNA amplicon sequencing and data analysis

To analyze cecal and colonic microbial composition, fecal samples were collected in sterile tubes, quickly frozen in liquid nitrogen, and stored at -80°C. Microbial DNA was extracted using the OMEGA Soil DNA Kit (Omega Bio-Tek, M5635-02, USA) according to the manufacturer’s instructions. DNA integrity was checked via agarose gel electrophoresis, and its concentration and purity were measured with a NanoDrop ND-2000 spectrophotometer (Thermo Scientific, USA). The hypervariable V3–V4 region of the 16S rRNA gene was amplified using specific primers 338F and 806R, followed by purification and sequencing with the Illumina MiSeq PE300 platform. (Illumina, San Diego, USA) [47].

After sequencing, raw reads were filtered for quality control using fastp [48] and merged with FLASH (V1.2.11) [49]. High-quality sequences were processed using DADA2 [50] to generate amplicon sequence variants (ASVs). The classification of ASVs is conducted using the Naive Bayes consensus taxonomy classifier implemented in QIIME2 (version 2020.2) [51].

Further bioinformatics analyses were conducted on the Majorbio Cloud platform. Based on the ASVs information, rarefaction curves and α-diversity indices were analyzed with Mothur v1.30.1 [52]. β-diversity was analyzed through principal coordinate analysis (PCoA) based on weighted UniFrac distances in the Vegan v2.5-3 package. The linear discriminant analysis (LDA) effect size (LEfSe) [49] (http://huttenhower.sph.harvard.edu/LEfSe) was performed to identify the differentially abundant taxa (phylum to genus) of bacteria. The correlation between colitis phenotype and microbes was performed by Spearman’s rank correlation.

### 16S rRNA Quantitative PCR

The total bacteria load in mice cecal and colonic samples was quantified by using 16S rRNA gene-based quantitative PCR assay as previously reported [53]. PCR amplification was conducted using bacterial 16S rRNA-targeting primers (forward: 5ʹ- AGAGTTTGATCCTGGCTCAG-3ʹ; reverse: 5ʹ-CTGCTGCCTYCCGTA-3ʹ). Quantitative PCR was performed on a Light Cycler® 480 II system (Roche, Switzerland) under the following thermal cycling conditions: an initial denaturation at 95°C for 5 min, followed by 45 cycles of 95°C for 10 s, 60°C for 10 s, and 72°C for 10 s. To ensure accurate quantification, a standard curve was established using a pMD18T vector incorporating a bacterial 16S rRNA-specific fragment for normalization. The sequences of the primers used are provided in Supplementary Table 3.

### Safety evaluation of Syn3

Eight-week-old male C57BL/6J mice were randomly divided into two groups. The mice were administrated with PBS or Syn3 for one month every three days. The body weight of the mice was recorded every three days. The food intake and water consumption of the mice were monitored every week. To investigate whether Syn3 treatment induced depression-like or anxiety-like behavior and affected the locomotor activity, we conducted the tail-suspension test on day 30. Mice were suspended by tail for 10 min, and the mobility/immobility time was recorded, and the analysis was performed by Noldus Etho Vision XT 11.5 [54]. For the pole-climbing test, the rubber ball was wrapped around the iron frame to avoid slippage. The mice were positioned with their heads straight upward and their tails down at the top of the pole. The locomotion activity time (T-LA) (i.e., when the hind feet of mice left the platform) and the turning time (T-Turn) were recorded. This experiment was conducted 9 times for each mouse, and the average values were calculated as the final results. The interval between each experiment was ≥15 min. All mice were trained three times per day for three consecutive days to become accustomed to the apparatus before the formal experiment [55]. On the 30th day of the experiment, the mice were euthanized, and the main organs (colon, cecum, spleen, kidney, liver, brain, thymus, testis) were collected to calculate the organ coefficients. The concentration of serum dopamine was assayed by ELISA (Sangon Biotech, D751019-0048,China). The activity of aspartate aminotransferase was assayed by a microplate assay kit (Nanjing Jiancheng, C010-2-1,China).

### Statistics

Statistical analysis data was presented as mean ± SEM. The comparisons between two groups of samples were evaluated using student’s *t*-test. Multiple comparison was performed by using one-way analysis of variance (ANOVA). The statistical significance was defined as: \**P* < 0.05, \*\**P* < 0.01, \*\*\**P* < 0.001, \*\*\*\**P* < 0.001. All statistical analyses were calculated using GraphPad Prism (10.1.2 software).

## Supporting information

all supplementary material for manuscript

## Author Contributions

P.J.Y performed the experiments, analyzed the results, and wrote the manuscript; W.J.Z and C.Y.L participated in article proofing. Q.S. and Y.P.Y. conceived and supervised the study. All authors contributed to manuscript revision, read and approved the submitted version.

## Acknowledgments

We acknowledge Prof. Sheng Yang of CAS Center for Excellence in Molecular Plant Sciences for kindly supplying the genome-editing vectors.

## Availability of data and material

The 16S rRNA amplicon sequencing data used in this study have been deposited in the Sequence Read Archive of NCBI (https://submit.ncbi.nlm.nih.gov/subs/) with Accession No. SUB14825764

## References

1. Wu G, Xu T, Zhao N, Lam YY, Ding X, Wei D, et al. A core microbiome signature as an indicator of health. Cell. 2024;187(23):6550–65 e11.

2. Sorboni SG, Moghaddam HS, Jafarzadeh-Esfehani R, Soleimanpour S. A Comprehensive Review on the Role of the Gut Microbiome in Human Neurological Disorders. Clin Microbiol Rev. 2022;35(1):e0033820.

3. Snigdha S, Ha K, Tsai P, Dinan TG, Bartos JD, Shahid M. Probiotics: Potential novel therapeutics for microbiota-gut-brain axis dysfunction across gender and lifespan. Pharmacol Ther. 2022;231:107978.

4. Effendi SSW, Ng IS. Prospective and challenges of live bacterial therapeutics from a superhero *Escherichia coli* Nissle 1917. Crit Rev Microbiol. 2023;49(5):611–27.

5. Chen H, Lei P, Ji H, Yang Q, Peng B, Ma J, et al. Advances in *Escherichia coli* Nissle 1917 as a customizable drug delivery system for disease treatment and diagnosis strategies. Mater Today Bio. 2023;18:100543.

6. Canale FP, Basso C, Antonini G, Perotti M, Li N, Sokolovska A, et al. Metabolic modulation of tumours with engineered bacteria for immunotherapy. Nature. 2021;598(7882):662-6.

7. Yu X, Lin C, Yu J, Qi Q, Wang Q. Bioengineered *Escherichia coli* Nissle 1917 for tumour-targeting therapy. Microb Biotechnol. 2020;13(3):629–36.

8. Zhao Z, Xu S, Zhang W, Wu D, Yang G. Probiotic *Escherichia coli* NISSLE 1917 for inflammatory bowel disease applications. Food Funct. 2022;13(11):5914–24.

9. Praveschotinunt P, Duraj-Thatte AM, Gelfat I, Bahl F, Chou DB, Joshi NS. Engineered *E. coli Nissle* 1917 for the delivery of matrix-tethered therapeutic domains to the gut. Nat Commun. 2019;10(1):5580.

10. Wu H, Wei J, Zhao X, Liu Y, Chen Z, Wei K, et al. Neuroprotective effects of an engineered *Escherichia coli* Nissle 1917 on Parkinson’s disease in mice by delivering GLP-1 and modulating gut microbiota. Bioeng Transl Med. 2023;8(5):e10351.

11. Yan X, Liu XY, Zhang D, Zhang YD, Li ZH, Liu X, et al. Construction of a sustainable 3-hydroxybutyrate-producing probiotic *Escherichia coli* for treatment of colitis. Cell Mol Immunol. 2021;18(10):2344–57.

12. Sun R, Yu P, Guo L, Huang Y, Nie Y, Yang Y. Improving the growth and intestinal colonization of *Escherichia coli* Nissle 1917 by strengthening its oligopeptides importation ability. Metab Eng. 2024;86:157–71.

13. Li J, Hou W, Lin S, Wang L, Pan C, Wu F, et al. Polydopamine Nanoparticle-Mediated Dopaminergic Immunoregulation in Colitis. Adv Sci (Weinh). 2022;9(1):e2104006.

14. Wang X, Cao Z, Zhang M, Meng L, Ming Z, Liu J. Bioinspired oral delivery of gut microbiota by self-coating with biofilms. Sci Adv. 2020;6(26):eabb1952.

15. Wang X, Gao S, Yun S, Zhang M, Peng L, Li Y, et al. Microencapsulating Alginate-Based Polymers for Probiotics Delivery Systems and Their Application. Pharmaceuticals (Basel). 2022;15(5).

16. Pochart P, Marteau P, Bouhnik Y, Goderel I, Bourlioux P, Rambaud JC. Survival of bifidobacteria ingested via fermented milk during their passage through the human small intestine: an in vivo study using intestinal perfusion. Am J Clin Nutr. 1992;55(1):78–80.

17. Li Z, Huang Z, Gu P. Response of *Escherichia coli* to Acid Stress: Mechanisms and Applications-A Narrative Review. Microorganisms. 2024;12(9).

18. Bender JK, Praszkier J, Wakefield MJ, Holt K, Tauschek M, Robins-Browne RM, et al. Involvement of PatE, a prophage-encoded AraC-like regulator, in the transcriptional activation of acid resistance pathways of enterohemorrhagic Escherichia coli strain EDL933. Appl Environ Microbiol. 2012;78(15):5083–92.

19. Zhao B, Houry WA. Acid stress response in enteropathogenic gammaproteobacteria: an aptitude for survival. Biochem Cell Biol. 2010;88(2):301–14.

20. Andreichin MA. [Antimicrobial properties of bile and bile acids]. Antibiotiki. 1980;25(12):936–9.

21. Paul S, Alegre KO, Holdsworth SR, Rice M, Brown JA, McVeigh P, et al. A single-component multidrug transporter of the major facilitator superfamily is part of a network that protects *Escherichia coli* from bile salt stress. Mol Microbiol. 2014;92(4):872–84.

22. Rodriguez-Beltran J, Rodriguez-Rojas A, Guelfo JR, Couce A, Blazquez J. The *Escherichia coli* SOS gene dinF protects against oxidative stress and bile salts. Plos One. 2012;7(4):e34791.

23. van Bokhorst-van de Veen H, Lee IC, Marco ML, Wels M, Bron PA, Kleerebezem M. Modulation of *Lactobacillus plantarum* gastrointestinal robustness by fermentation conditions enables identification of bacterial robustness markers. Plos One. 2012;7(7):e39053.

24. Troge A, Scheppach W, Schroeder BO, Rund SA, Heuner K, Wehkamp J, et al. More than a marine propeller--the flagellum of the probiotic *Escherichia coli* strain Nissle 1917 is the major adhesin mediating binding to human mucus. Int J Med Microbiol. 2012;302(7-8):304–14.

25. Hammar M, Arnqvist A, Bian Z, Olsen A, Normark S. Expression of two csg operons is required for production of fibronectin-and congo red-binding curli polymers in *Escherichia coli* K-12. Mol Microbiol. 1995;18(4):661–70.

26. Frenzel E, Legebeke J, van Stralen A, van Kranenburg R, Kuipers OP. In vivo selection of sfGFP variants with improved and reliable functionality in industrially important thermophilic bacteria. Biotechnol Biofuels. 2018;11:8.

27. Scaldaferri F, Gerardi V, Mangiola F, Lopetuso LR, Pizzoferrato M, Petito V, et al. Role and mechanisms of action of *Escherichia coli* Nissle 1917 in the maintenance of remission in ulcerative colitis patients: An update. World J Gastroenterol. 2016;22(24):5505–11.

28. Vallejos OP, Bueno SM, Kalergis AM. Probiotics in inflammatory bowel disease: microbial modulation and therapeutic prospects. Trends Mol Med. 2025.

29. Wu Z, Huang S, Li T, Li N, Han D, Zhang B, et al. Gut microbiota from green tea polyphenol-dosed mice improves intestinal epithelial homeostasis and ameliorates experimental colitis. Microbiome. 2021;9(1):184.

30. Joo M, Nam S. Adolescent gut microbiome imbalance and its association with immune response in inflammatory bowel diseases and obesity. BMC Microbiol. 2024;24(1):268.

31. Ma Y, Fu W, Hong B, Wang X, Jiang S, Wang J. Antibacterial MccM as the Major Microcin in *Escherichia coli Nissle* 1917 against Pathogenic Enterobacteria. Int J Mol Sci. 2023;24(14).

32. Fang K, Jin X, Hong SH. Probiotic *Escherichia coli* inhibits biofilm formation of pathogenic E. coli via extracellular activity of DegP. Sci Rep. 2018;8(1):4939.

33. Kleta S, Nordhoff M, Tedin K, Wieler LH, Kolenda R, Oswald S, et al. Role of F1C fimbriae, flagella, and secreted bacterial components in the inhibitory effect of probiotic Escherichia coli Nissle 1917 on atypical enteropathogenic *E. coli* infection. Infect Immun. 2014;82(5):1801–12.

34. Trebichavsky I, Splichal I, Rada V, Splichalova A. Modulation of natural immunity in the gut by *Escherichia coli* strain Nissle 1917. Nutr Rev. 2010;68(8):459–64.

35. Garcia Mansilla MJ, Rodriguez Sojo MJ, Lista AR, Ayala Mosqueda CV, Ruiz Malagon AJ, Galvez J, et al. Exploring Gut Microbiota Imbalance in Irritable Bowel Syndrome: Potential Therapeutic Effects of Probiotics and Their Metabolites. Nutrients. 2024;17(1).

36. Hu R, Lin H, Li J, Zhao Y, Wang M, Sun X, et al. Probiotic *Escherichia coli* Nissle 1917-derived outer membrane vesicles enhance immunomodulation and antimicrobial activity in RAW264.7 macrophages. BMC Microbiol. 2020;20(1):268.

37. Li S, Zhuge A, Wang K, Lv L, Bian X, Yang L, et al. Ketogenic diet aggravates colitis, impairs intestinal barrier and alters gut microbiota and metabolism in DSS-induced mice. Food Funct. 2021;12(20):10210–25.

38. Shin NR, Whon TW, Bae JW. Proteobacteria: microbial signature of dysbiosis in gut microbiota. Trends Biotechnol. 2015;33(9):496–503.

39. Luo J, Wang Z, Fan B, Wang L, Liu M, An Z, et al. A comparative study of the effects of different fucoidans on cefoperazone-induced gut microbiota disturbance and intestinal inflammation. Food Funct. 2021;12(19):9087–97.

40. Yang Y, Lang N, Yang G, Yang S, Jiang W, Gu Y. Improving the performance of *solventogenic clostridia* by reinforcing the biotin synthetic pathway. Metab Eng. 2016;35:121–8.

41. Jiang Y, Chen B, Duan C, Sun B, Yang J, Yang S. Multigene editing in the *Escherichia coli* genome via the CRISPR-Cas9 system. Appl Environ Microbiol. 2015;81(7):2506–14.

42. Tummala SB, Welker NE, Papoutsakis ET. Development and characterization of a gene expression reporter system for *Clostridium acetobutylicum* ATCC 824. Appl Environ Microbiol. 1999;65(9):3793–9.

43. Castanie-Cornet MP, Penfound TA, Smith D, Elliott JF, Foster JW. Control of acid resistance in *Escherichia coli*. J Bacteriol. 1999;181(11):3525–35.

44. Vogel HJ, Bonner DM. Acetylornithinase of *Escherichia coli*: partial purification and some properties. J Biol Chem. 1956;218(1):97–106.

45. Ren C, Gu Y, Wu Y, Zhang W, Yang C, Yang S, et al. Pleiotropic functions of catabolite control protein CcpA in Butanol-producing *Clostridium acetobutylicum*. BMC Genomics. 2012;13:349.

46. Zhou J, Li M, Chen Q, Li X, Chen L, Dong Z, et al. Programmable probiotics modulate inflammation and gut microbiota for inflammatory bowel disease treatment after effective oral delivery. Nat Commun. 2022;13(1):3432.

47. Guo L, Xu L, Nie Y, Liu L, Liu Z, Yang Y. Murine gut microbial interactions exert antihyperglycemic effects. ISME J. 2025;19(1).

48. Chen S, Zhou Y, Chen Y, Gu J. fastp: an ultra-fast all-in-one FASTQ preprocessor. Bioinformatics. 2018;34(17):i884–i90.

49. Segata N, Izard J, Waldron L, Gevers D, Miropolsky L, Garrett WS, et al. Metagenomic biomarker discovery and explanation. Genome Biol. 2011;12(6):R60.

50. Callahan BJ, McMurdie PJ, Rosen MJ, Han AW, Johnson AJ, Holmes SP. DADA2: High-resolution sample inference from Illumina amplicon data. Nat Methods. 2016;13(7):581–3.

51. Bolyen E, Rideout JR, Dillon MR, Bokulich NA, Abnet CC, Al-Ghalith GA, et al. Reproducible, interactive, scalable and extensible microbiome data science using QIIME 2. Nat Biotechnol. 2019;37(8):852–7.

52. Schloss PD, Westcott SL, Ryabin T, Hall JR, Hartmann M, Hollister EB, et al. Introducing mothur: open-source, platform-independent, community-supported software for describing and comparing microbial communities. Appl Environ Microbiol. 2009;75(23):7537–41.

53. Yang Y, Yu P, Lu Y, Gao C, Sun Q. Disturbed rhythmicity of intestinal hydrogen peroxide alters gut microbial oscillations in BMAL1-deficient monkeys. Cell Rep. 2023;42(3):112183.

54. Yao C, Jiang N, Sun X, Zhang Y, Pan R, He Q, et al. Effects of inulin-type oligosaccharides (JSO) from Cichorium intybus L. on behavioral deficits induced by chronic restraint stress in mice and associated molecular alterations. Front Pharmacol. 2024;15:1484337.

55. Xu YD, Cui C, Sun MF, Zhu YL, Chu M, Shi YW, et al. Neuroprotective Effects of Loganin on MPTP-Induced Parkinson’s Disease Mice: Neurochemistry, Glial Reaction and Autophagy Studies. J Cell Biochem. 2017;118(10):3495–510.

