## Supplementary material for "Construction of an engineered *Escherichia coli* with enhanced intestinal colonization and anti-inflammatory efficacy in colitis": all supplementary material for manuscript

**Fig. S1 LacZ reporter-based screening of strong promoters that suitable for EcN under the aerobic and anaerobic conditions.** The plasmids containing the *LacZ* reporter that driven by different promoters ($P_{less}$, $P_{adc}$, Plp$_{_0237}$, $P_{CAC3456}$, Plp$_{_0775}$, $P_{sol}$, and $P_{bio}$) were electroporated into EcN. The plasmid-containing strains were inoculated into M9 medium. The $OD_{600}$ and β-galactosidase activity were assayed at 12 h under aerobic (**a**) and anaerobic (**b**) conditions.

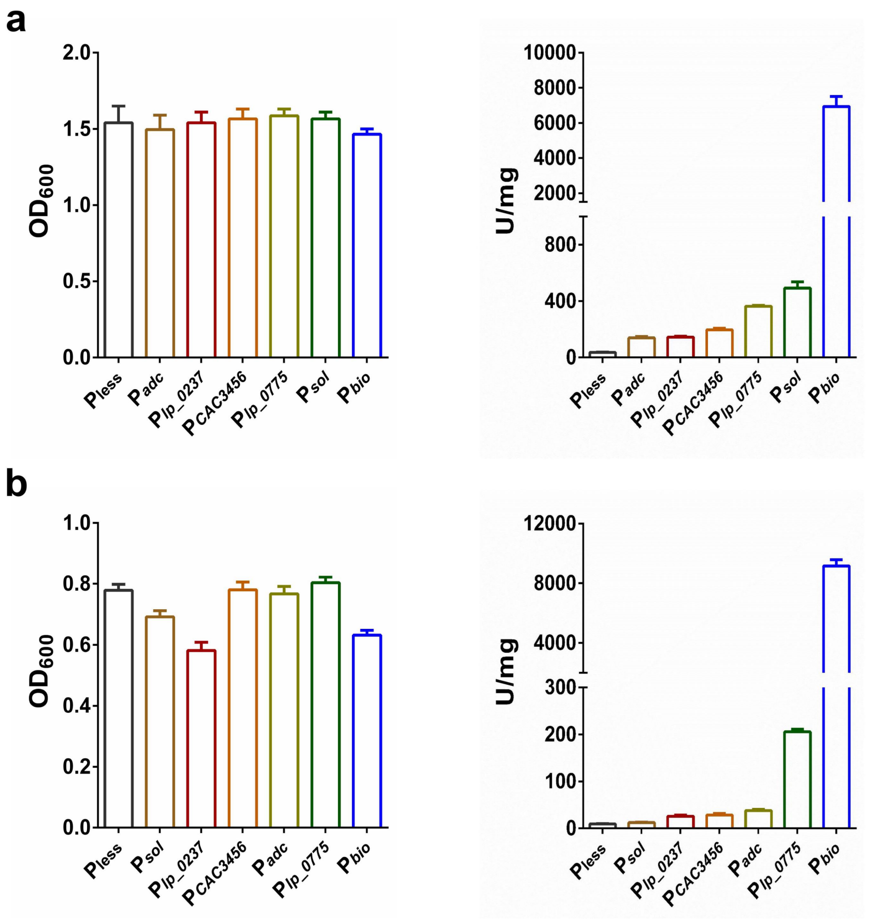

**Fig. S2 The growth curve of engineered EcN strains. a** The growth curve of EcN-P and EcN-PatE. **b** The growth curve of EcN-P, EcN-MdtM, and EcN-DinF.

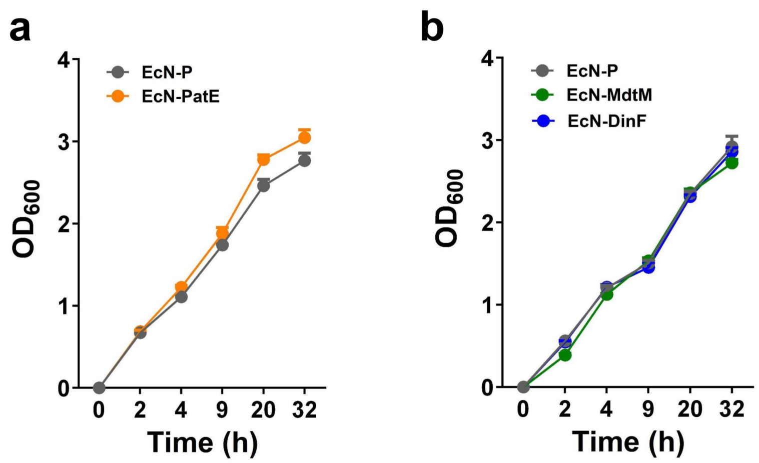

**Fig. S3. PCR-based verification of genome integration of P$_{bio}$-PatE-MdtM in EcN-PM and Syn3.** The markers are labelled as bended arrows.

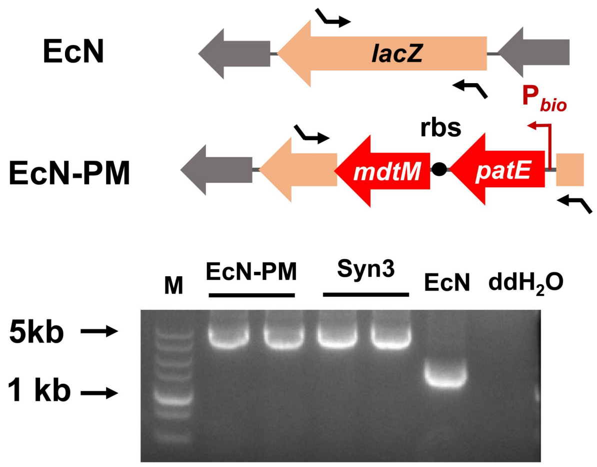

**Fig. S4 The gut microbial alterations of colitis mice after Syn3 administration.** The average Good's coverage of each sample in the cecal (**a**) and colonic (**b**) microbiome of the NC, PBS, EcN, and Syn3 groups. The cecal (**c**) and colonic (**d**) microbial dysbiosis index of four mice groups. LEfSe analysis-based characterization of the taxonomic differences of cecal (**e**) and colonic (**f**) microbiome among the four mice groups. LDA score cut-off was set as 2.0 (***$P < 0.001$, ****P< 0.0001)

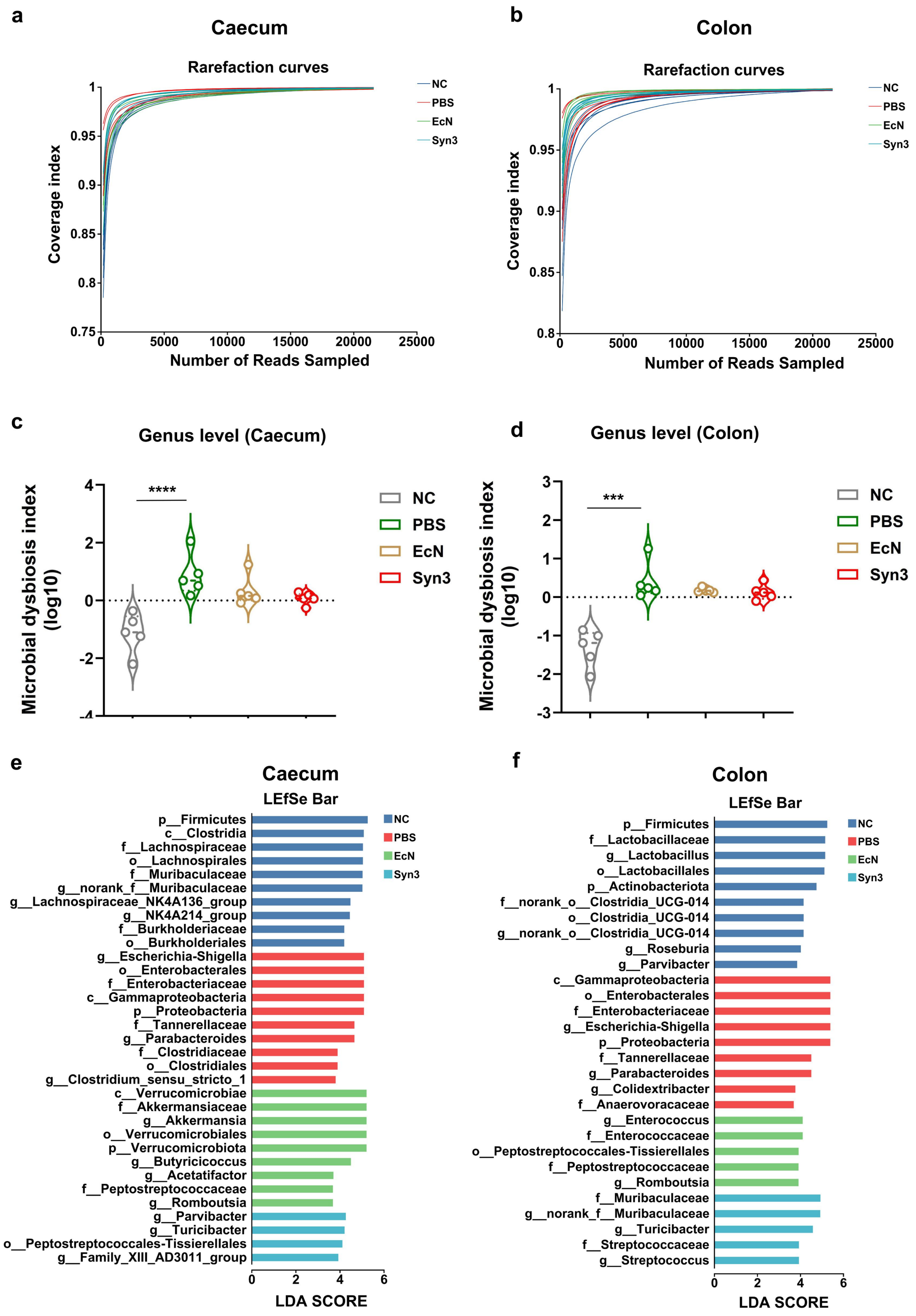
a
Caecum
Rarefaction curves
Coverage index
Number of Reads Sampled
NC
PBS
EcN
Syn3
b
Colon
Rarefaction curves
Coverage index
Number of Reads Sampled
c
Genus level (Caecum)
Microbial dysbiosis index (log10)
****
d
Genus level (Colon)
Microbial dysbiosis index (log10)
***
e
Caecum
LEfSe Bar
p__Firmicutes
c__Clostridia
f__Lachnospiraceae
o__Lachnospirales
f__Muribaculaceae
g__norank_f__Muribaculaceae
g__Lachnospiraceae_NK4A136_group
g__NK4A214_group
f__Burkholderiaceae
o__Burkholderiales
g__Escherichia-Shigella
o__Enterobacterales
f__Enterobacteriaceae
c__Gammaproteobacteria
p__Proteobacteria
f__Tannerellaceae
g__Parabacteroides
f__Clostridiaceae
o__Clostridiales
g__Clostridium_sensu_stricto_1
c__Verrucomicrobiae
f__Akkermansiaceae
g__Akkermansia
o__Verrucomicrobiales
p__Verrucomicrobiota
g__Butyricicoccus
g__Acetatifactor
f__Peptostreptococcaceae
g__Romboutsia
g__Parvibacter
g__Turicibacter
o__Peptostreptococcales-Tissierellales
g__Family_XIII_AD3011_group
LDA SCORE
f
Colon
LEfSe Bar
p__Firmicutes
f__Lactobacillaceae
g__Lactobacillus
o__Lactobacillales
p__Actinobacteriota
f__norank_o__Clostridia_UCG-014
o__Clostridia_UCG-014
g__norank_o__Clostridia_UCG-014
g__Roseburia
g__Parvibacter
c__Gammaproteobacteria
o__Enterobacterales
f__Enterobacteriaceae
g__Escherichia-Shigella
p__Proteobacteria
f__Tannerellaceae
g__Parabacteroides
g__Colidextribacter
f__Anaerovoracaceae
g__Enterococcus
f__Enterococcaceae
o__Peptostreptococcales-Tissierellales
f__Peptostreptococcaceae
g__Romboutsia
f__Muribaculaceae
g__norank_f__Muribaculaceae
g__Turicibacter
f__Streptococcaceae
g__Streptococcus
LDA SCORE

**Fig. S5 The total bacterial load in cecal and colonic samples of four mice groups.**

**a** Comparison of total bacterial load in cecal samples among the four mice groups. **b** Comparison of total bacterial load in colonic samples among the four mice groups. Data is presented in mean ± SEM.

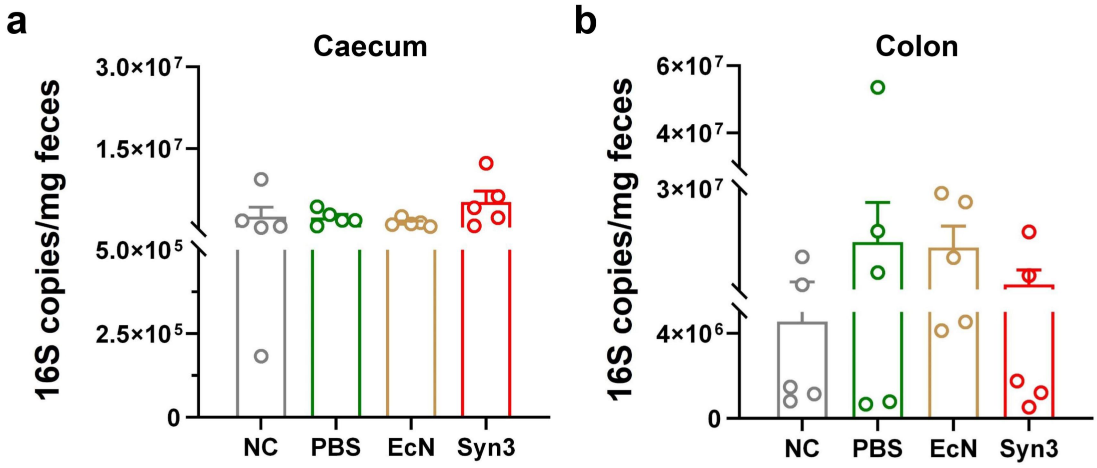

**Fig. S6. *In vivo* evaluation of biosafety of Syn3. a** Schematic illustration of animal experiment. **b** Comparison of weight and weight change between the PBS and Syn3 gavaged mice. **c** Comparison of the food and water intake between the PBS and Syn3 gavaged mice. **d** The concentration of dopamine in the serum of two mice groups. **e** The percentage of immobility and mobility time in the tail-hanging test. **f** Time to descend of mice in pole-climbing test. **g** Comparison of classical immune cells level between the two mice groups. **h** Comparison of HGB and MCHC levels between the two mice groups. **i** Comparison of the glutamate oxaloacetate transaminase (GOT) activity between the two mice groups. **j** The length of cecum and colon in two mice groups. **k** The main organ coefficient (organ weight/body weight) of two mice groups. The statistical significance between two groups was analyzed using *t*-test (*$P$ <0.05).

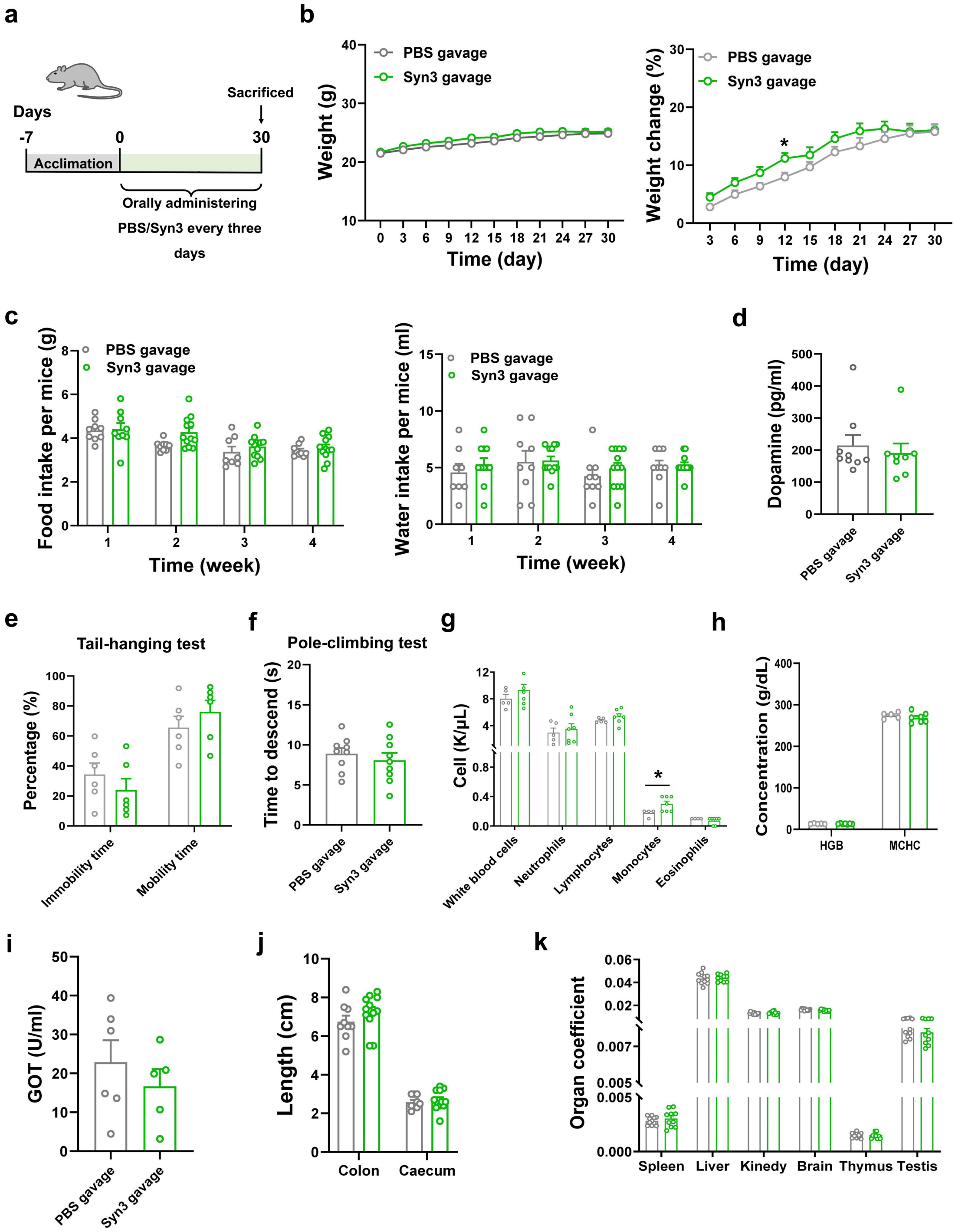
a
Days
-7
0
30
Sacrificed
Acclimation
Orally administering PBS/Syn3 every three days
b
PBS gavage
Syn3 gavage
Weight (g)
Time (day)
Weight change (%)
Time (day)
c
Food intake per mice (g)
Time (week)
Water intake per mice (ml)
Time (week)
d
Dopamine (pg/ml)
PBS gavage
Syn3 gavage
e
Tail-hanging test
Percentage (%)
Immobility time
Mobility time
f
Pole-climbing test
Time to descend (s)
g
Cell (K/μL)
White blood cells
Neutrophils
Lymphocytes
Monocytes
Eosinophils
h
Concentration (g/dL)
HGB
MCHC
i
GOT (U/ml)
j
Length (cm)
Colon
Caecum
k
Organ coefficient
Spleen
Liver
Kinedy
Brain
Thymus
Testis

**Table S1. Strains used in this study.**

| Strains | Description | Source |
|---|---|---|
| *Escherchia coli* Nissle 1917 | Non-pathogenic probiotic strain, pMUT1, pMUT2, microcin H47/M, F1C-fimbria, type 1 fimbria, curli | Pharma-Zentrale GmbH |
| Top10 | For general gene cloning and plasmid construction | Invitrogen, C404003 |
| EcN-P | EcN, carries the pET-32a-$P_{bio}$ plasmid | This study |
| EcN-PatE | EcN, carries the pET-32a-$P_{bio}$-*patE* plasmid | This study |
| EcN-MdtM | EcN, carries the pET-32a-$P_{bio}$-*mdtM* plasmid | This study |
| EcN-DinF | EcN, carries the pET-32a-$P_{bio}$-*dinF* plasmid | This study |
| EcN-PatE-MdtM | EcN, carries the pET-32a-$P_{bio}$-*patE-mdtM* plasmid | This study |
| EcN-PM | EcN, inserting the *patE* and *mdtM* into *lacZ* site | This study |
| EcN-Csg | EcN, deleting the *csgD*, and replacing the promoters of curli fiber formation genes | This study |
| Syn3 | EcN, inserting the $P_{bio}$-*patE-mdtM* into *lacZ* site, deleting the *csgD*, and replacing the promoters of curli fiber formation genes | This study |

| | | |
|---|---|---|
| EcN-fluo | EcN, carrying pTD103luxI_sfGFP | This study |
| EcN-PM-fluo | EcN-PM, carrying pTD103luxI_sfGFP | This study |
| EcN-Csg-fluo | EcN-Csg, carrying pTD103luxI_sfGFP | This study |
| Syn3-fluo | Syn3, carrying pTD103luxI_sfGFP | This study |

**Table S2. Plasmids used in this study.**

| Plasmid | Description | Source |
|---|---|---|
| pET-32a-$P_{bio}$ | the control vector | This study |
| pET-32a-$P_{bio}$-*patE* | *patE* overexpression vector, derived from pET-32a-$P_{bio}$ | This study |
| pET-32a-$P_{bio}$-*mdtM* | *mdtM* overexpression vector, derived from pET-32a- $P_{bio}$ | This study |
| pET-32a-$P_{bio}$-*dinF* | *dinF* overexpression vector, derived from pET-32a- $P_{bio}$ | This study |
| pET-32a-$P_{bio}$-*patE-mdtM* | *patE* and *mdtM* overexpression vector, derived from pET-32a-$P_{bio}$-*patE* | This study |
| pCas | RepA101(Ts) kan Pcas-cas9 ParaB-Red lacIq Ptrc-sgRNA-pMB1 | [1] |
| pTargetF | Plasmid harboring sgRNAs without donor DNAs | [1] |
| pET-32a-*lacZ* | Vector used for β-Galactosidase assay, | This study |
| pET-32a-$P_{adc}$-*lacZ* | lacZ reporter gene driven by $P_{adc}$ | This study |
| pET-32a-$P_{sol}$-*lacZ* | lacZ reporter gene driven by $P_{sol}$ | This study |
| pET-32a-$P_{bio}$-*lacZ* | lacZ reporter gene driven by $P_{bio}$ | This study |
| pET-32a-$P_{CAC3456}$-*lacZ* | lacZ reporter gene driven by $P_{CAC3456}$ | This study |
| pET-32a-$P_{lp\_0237}$-*lacZ* | lacZ reporter gene driven by $P_{lp\_0237}$ | This study |
| pET-32a-$P_{lp\_0775}$-*lacZ* | lacZ reporter gene driven by $P_{lp\_0775}$ | This study |
| pTD103luxI_sfGFP | Express sfGFP fluorescence for *in vivo* imaging experiments (IVIS) | Ningbo Naisi technology Co., Ltd. |
| pTargetF-lacZ-sg1-$P_{bio}$-*patE-mdtM* | The plasmid used for inserting $P_{bio}$-*patE-mdtM* into lacZ site | This study |
| pTargetF-CsgD sg1-$P_{bio}$-Plp$_{_0775}$ | pTarget harboring sgRNA targeting *csgD*, the promoter of two Csg operon will be replaced with $P_{bio}$ and $P_{lp\_0775}$. | This study |

**Table S3. Primers used in this study.**

| Primer | Sequence (5'→3') | Description |
|---|---|---|
| **Quantitative PCR assay** | | |
| qRT-*gadA*-s | CTACTATCGCGGAGTCAAAACG | Forward qRT-PCR primer for *gadA* (CIW80_12250) |
| qRT-*gadA*-a | ATGCCACAGATCGGCAACCATG | Reverse qRT-PCR primer for *gadA* (CIW80_12250) |
| qRT-*gadB*-s | CGGAACTACTCGATTCACGTTTTG | Forward qRT-PCR primer for *gadB* |
| qRT-*gadB*-a | GCTGCAGATTGCGGATATTCTTC | Reverse qRT-PCR primer for *gadB* |
| qRT-*gadE*-s | AAATCAATTCCCTGTCAGAGATCA | Forward qRT-PCR primer for *gadE* |
| qRT-*gadE*-a | GGCAAGTGTTTACCATAAGTAATC | Reverse qRT-PCR primer for *gadE* |
| qRT-*gadW*-s | CGAGAGTATCCTGCTGCTGGAT | Forward qRT-PCR primer for *gadW* |
| qRT-*gadW*-a | AGCATTTTGCTCAGTATCGGCG | Reverse qRT-PCR primer for *gadW* |
| qRT-*gadX*-s | GCTATTTTAATGGCGGTGATCTG | Forward qRT-PCR primer for *gadX* |
| qRT-*gadX*-a | CTGCGATAGTTGCGCAACTTCC | Reverse qRT-PCR primer for *gadX* |
| qRT-*hdeA*-s | CGTTATTCTTGGTGGTCTGCTTC | Forward qRT-PCR primer for *hdeA* |
| qRT-*hdeA*-a | GGTTGCAATACCCTGAACATCT | Reverse qRT-PCR primer for *hdeA* |
| qRT-*hdeD*-s | GCAGAGCAATCCAGTTTATTGC | Forward qRT-PCR primer for *hdeD* |

| | | |
|---|---|---|
| qRT-16s rRNA-s | AACACATGCAAGTCGAGGGG | Forward qRT-PCR primer for 16S rRNA |
| qRT-16s rRNA-a | CAGACGCATCCCCATCCATC | Reverse qRT-PCR primer for 16S rRNA |
| qRT-*patE*-s | CGGTGATCCTCGTTCGTCG | Forward qRT-PCR primer for *patE* |
| qRT-*patE*-a | AATTGCGTGGCAACTGACG | Reverse qRT-PCR primer for *patE* |
| qRT-*mdtM*-s | GGCGACTCTACTTGCCGAGC | Forward qRT-PCR primer for *mdtM* |
| qRT-*mdtM*-a | CTGGGCTTTACGTTGTGCGA | Reverse qRT-PCR primer for *mdtM* |
| qRT-*csgB*-s | TGCTCAGTTACGGCAGGGAG | Forward qRT-PCR primer for *csgB* |
| qRT-*csgB*-a | TGGCATCGTTGGCACTGCCC | Reverse qRT-PCR primer for *csgB* |
| qRT-*csgD*-s | AGGAAACCGCTTGTGTCCGG | Forward qRT-PCR primer for *csgD* |
| qRT-*csgD*-a | CAGCCCTCCTTACTCATCGG | Reverse qRT-PCR primer for *csgD* |
| qRT-*csgE*-s | TTGTGGCGGCAGAATTTCTG | Forward qRT-PCR primer for *csgE* |
| qRT-*csgE*-a | CCCGTATAGTCACTTTCCC | Reverse qRT-PCR primer for *csgE* |
| **16S rRNA gene amplicon sequencing** | | |
| 338F | ACTCCTACGGGAGGCAGCA | Forward primer for the V3-V4 region of 16S rRNA |
| 806R | GGACTACHVGGGTWTCTAAT | Reverse primer for the V3-V4 |

| | | |
|---|---|---|
| | | region of 16S rRNA |

**REFERENCES**

1. Jiang Y, Chen B, Duan C, Sun B, Yang J, Yang S. Multigene editing in the Escherichia coli genome via the CRISPR-Cas9 system. Appl Environ Microbiol. 2015;81(7):2506-14.
